# An integrative genomic analysis of the Longshanks selection experiment for longer limbs in mice

**DOI:** 10.1101/378711

**Authors:** João P. L. Castro, Michelle N. Yancoskie, Marta Marchini, Stefanie Belohlavy, Marek Kučka, William H. Beluch, Ronald Naumann, Isabella Skuplik, John Cobb, Nick H. Barton, Campbell Rolian, Yingguang Frank Chan

**Author notes:** **Corresponding author:** Campbell Rolian, Yingguang Frank Chan. indicates equal contribution (João P. L. Castro, Michelle N. Yancoskie). indicates equal contribution (Campbell Rolian, Yingguang Frank Chan).

## Abstract

Evolutionary studies are often limited by missing data that are critical to understanding the history of selection. Selection experiments, which reproduce rapid evolution under controlled conditions, are excellent tools to study how genomes evolve under strong selection. Here we present a genomic dissection of the Longshanks selection experiment, in which mice were selectively bred over 20 generations for longer tibiae relative to body mass, resulting in 13% longer tibiae in two replicate lines. We synthesized evolutionary theory, genome sequences and molecular genetics to understand the selection response and found that it involved both polygenic adaptation and discrete loci of major effect, with the strongest loci likely to be selected in parallel between replicates. We show that selection may favor de-repression of bone growth through inactivation of two limb enhancers of an inhibitor, *Nkx3-2*. Our integrative genomic analyses thus show that it is possible to connect individual base-pair changes to the overall selection response.

## Main Text

Understanding how populations adapt to a changing environment is an urgent challenge of global significance. The problem is especially acute for mammals, which often feature small and fragmented populations due to widespread habitat loss. Small populations often have limited capacity for rapid adaptation due to loss of diversity through inbreeding (*1*). From a genetic perspective, adaptation depends on the availability of beneficial alleles, which in mammals typically come from standing genetic variation, rather than new mutations (as in clonal microbes). Mammals nonetheless respond readily to selection in many traits, both in nature and in the laboratory (*2*-*6*). This is interpreted in quantitative genetics as evidence for a large set of nearly neutral loci; data from selection experiments are largely indistinguishable from this infinitesimal model, which is the basis for commercial breeding (*7*). However, it remains unclear what type of genomic change is associated with rapid response to selection, especially in small populations where allele frequency changes can be dominated by random drift. While a large body of theory exists to describe the birth, rise and eventual fixation of adaptive variants under diverse selection scenarios (*8*-*13*), few empirical datasets capture sufficient detail on the founding conditions and selection regime to allow full reconstruction of the selection response. This is particularly problematic in nature, where historical samples, environmental measurements and replicates are often missing. Selection experiments, which reproduce rapid evolution under controlled conditions, are therefore excellent tools to understand response to selection—and by extension—adaptive evolution in nature (4).

Here we describe an integrative, multi-faceted investigation into an artificial selection experiment, called Longshanks, in which mice were selected for increased tibia length, relative to body mass (*14*). The mammalian limb is an ideal model to study the dynamics of complex traits under selection. It is both morphologically complex and functionally diverse, reflecting its adaptive value; limb development has been studied extensively in mammals, birds and fishes as a genetic and evolutionary paradigm (*15*). The Longshanks selection experiment thus offers the opportunity to study selection response not only from a quantitative and population genetics perspective, but also from a developmental (*16*) and genomic perspective.

By design, the Longshanks experiment preserves a nearly complete archive of the phenotype (trait measurements) and genotype (via tissue samples) in the pedigree. By sequencing the initial and final genomes, we can study this example of rapid evolution with unprecedented detail and resolution. Here, with essentially complete information, we wish to answer a number of important questions regarding the factors that determine and constrain rapid adaptation. We ask whether the observed changes in gene frequency are due to selection, or random drift: does rapid selection response of a complex trait proceed through innumerable loci of infinitesimally small effect, or through a few loci of large effect? If so, what signature of selection do we expect? Finally, when the same trait changes occur independently, do these depend on changes in the same gene(s) or the same pathways (parallelism)?

### Longshanks selection for longer tibiae

At the start of the Longshanks experiment, we established three base populations with 14 pairs each by sampling from a genetically diverse, commercial mouse stock [Hsd:ICR, also known as CD-1; derived from mixed breeding of classical laboratory mice (*17*)]. In two replicate “Longshanks” lines (LS1 and LS2), we selectively bred mice with the longest tibia relative to the cube root of body mass, 15–20% of offspring being selected for breeding [see (*14*) for details]. We kept a third Control line (Ctrl) using an identical breeding scheme, except that breeders were selected at random. In LS1 and LS2, we observed a strong and significant response to selection of tibia length [0.29 and 0.26 Haldane or standard deviations (s.d.) per generation, from a selection differential of 0.73 s.d. in LS1 and 0.62 s.d. in LS2]. Over 20 generations, selection for longer relative tibia length produced increases of 5.27 and 4.81 s.d. in LS1 and LS2, respectively (or 12.7% and 13.1% in tibia length), with negligible changes in body mass (Fig. 1B & C; fig. S1). By contrast, Ctrl showed no directional change in tibia length or body mass (Fig. 1C; Student’s *t*-test, *P > 0.05*). This approximately 5 s.d. change in 20 generations is rapid compared to typical rates observed in nature [(*18*), but see (*19*)] but is in line with responses seen in selection experiments (*3*, *6*, *20*, *21*).

**Fig. 1.**
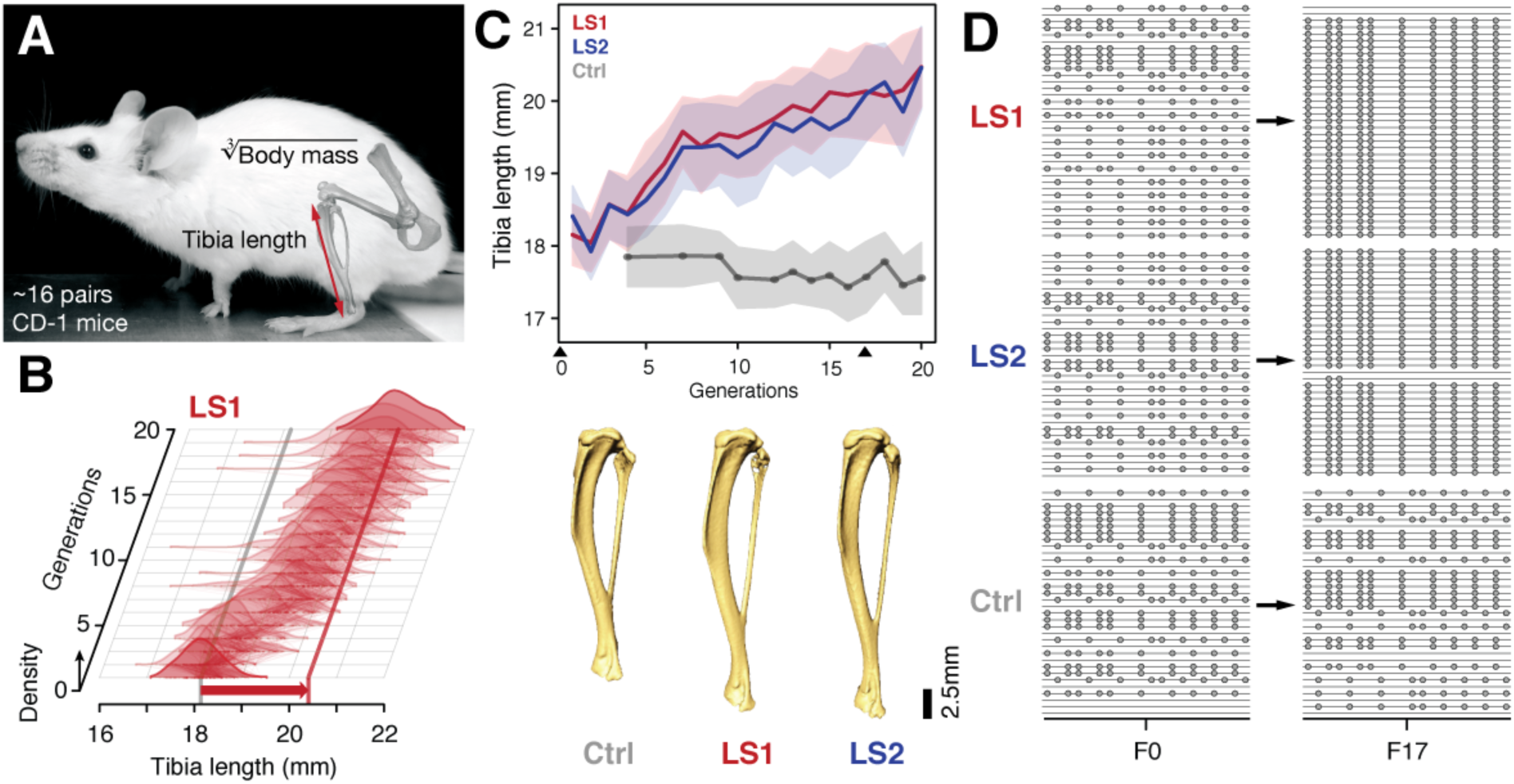
Selection for Longshanks mice produced rapid increase in tibia length. (**A** and **B**) Tibia length varies as a quantitative trait among outbred mice derived from the Hsd:ICR (also known as CD-1) commercial stock. Selective breeding for mice with the longest tibiae relative to body mass within families has produced a strong selection response in tibia length over 20 generations in Longshanks mice (13%, red arrow, LS1). (**C**) Both LS1 and LS2 produced replicated rapid increase in tibia length (red and blue; line and shading show mean ± s.d.) compared to random-bred Controls (grey). Arrowheads mark sequenced generations F0 and F17. See fig. S1 for body mass data. Lower panel: Representative tibiae from the Ctrl, LS1 and LS2 after 20 generations of selection. (**D**) Analysis of sequence diversity data (linked variants or haplotypes: lines; variants: dots) may detect signatures of selection, such as selective sweeps (F17 in LS1 and LS2) that result from selection favoring a particular variant (dots), compared to neutral or background patterns (Ctrl). Alternatively, selection may elicit a polygenic response, which may involve minor shifts in allele frequency at many loci and therefore may leave very different selection signature as shown here.

### Simulating selection response: infinitesimal model with linkage

We cannot use a neutral null model, since we know that strong selection was applied. We therefore developed a simulation that faithfully recapitulates the artificial selection experiment by integrating the trait measurements, selection regime, pedigree and genetic diversity of the Longshanks selection experiment, in order to generate an accurate expectation for the genomic response. Using the actual pedigree and trait measurements, we mapped fitness onto tibia length ***T*** and body mass ***B*** as a single composite trait *ln*(***TB*^*θ*^**). We estimated θ from actual data as −0.57, such that the ranking of breeders closely matched the actual composite ranking used to select breeders in the selection experiment, based on ***T*** and 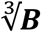 separately (*14*) (fig. S2A). We assumed a maximally polygenic genetic architecture using an “infinitesimal model with linkage” (abbreviated here as *HINF*), under which the trait is controlled by very many loci, each of infinitesimally small effect (see Supplementary Notes for details). Results from simulations seeded with actual genotypes or haplotypes showed that overall, the predicted increase in inbreeding closely matched the observed data (fig. S2B). We tested models with varying selection intensity and initial linkage disequilibrium (LD), and for each, ran 100 simulated replicates to determine the significance of changes in allele frequency (fig. S2C–E). This flexible quantitative genetics framework allows us to explore possible changes in genetic diversity over 17 generations of breeding, under strong selection.

In simulations, we followed blocks of genome as they were passed down the pedigree. In order to compare with observations, we seeded the initial genomes with single nucleotide polymorphisms (SNPs) in the same number and initial frequencies as the data. We observed much more variation between chromosomes in overall inbreeding (fig. S2B) and in the distribution of allele frequencies (fig. S3B) than expected from simulations in which the ancestral SNPs were initially in linkage equilibrium. This can be explained by linkage disequilibrium (LD) between the ancestral SNPs, which greatly increases random variation. Therefore, we based our significance tests on simulations that were seeded with SNPs drawn with LD consistent with the initial haplotypes (fig. S2C & E; see Supplementary Notes).

Because our simulations assume infinitesimal effects of loci, allele frequency shifts exceeding this stringent threshold suggest that discrete loci contribute significantly to the selection response. An excess of such loci in either a single LS replicate or in parallel would thus imply a mixed genetic architecture of a few large effect loci amid an infinitesimal background.

### Sequencing the Longshanks mice reveals genomic signatures of selection

To detect the genomic changes in the actual Longshanks experiment, we sequenced all individuals of the founder (F0) and 17^th^ generation (F17) to an average of 2.91-fold coverage (range: 0.73–20.6×; n = 169 with <10% missing F0 individuals; Table S1). Across the three lines, we found similar levels of diversity, with an average of 6.7 million (M) segregating SNPs (approximately 0.025%, or 1 SNP per 4 kbp; Table S1; figs S3A & S4). We checked the founder populations to confirm negligible divergence between the three founder populations (across-line F_ST_ on the order of 1×10^-4^), which increased to 0.18 at F17 (Table S2). This is consistent with random sampling from an outbred breeding stock. By F17, the number of segregating SNPs dropped to around 5.8 M (Table S1). This 13% drop in diversity (0.9M SNPs genome-wide) closely matched our simulation results, from which we could determine that the drop was mostly due to inbreeding, with only a minor contribution from selection (Supplementary Notes, fig. S2B, D).

We conclude that despite the strong selection on the LS lines, there was little perturbation to genome-wide diversity. Indeed, the changes in diversity during the 17 generations were remarkably similar in all three lines, despite Ctrl not having experienced selection on relative tibia length (fig. S3A). Hence, and consistent with our simulation results (fig. S2B, D), changes in global genome diversity had little power to distinguish selection from neutral drift despite the strong *phenotypic* selection response.

We next asked whether specific loci reveal more definitive differences between the LS replicates and Ctrl (and from infinitesimal predictions). We calculated Δz^2^, the square of arc-sin transformed allele frequency difference between F0 and F17; this has an expected variance of 1/2*N*_*e*_ per generation, independent of starting frequency, and ranges from 0 to π^2^. We averaged Δz^2^ within 10 kbp windows (see Methods for details), and found 169 windows belonging to eight clusters that had significant shifts in allele frequency in LS1 and/or 2 (corresponding to 9 and 164 clustered windows respectively at *P* ≤ 0.05 under *H*_*INF, max LD*_; Δz^2^ ≥ 0.33 π^2^; genome-wide Δz^2^= 0.02 ± 0.03 π^2^; Fig. 2; figs. S2D, S5, S6; see Methods for details) and in 3 clusters in Ctrl (8 windows; genome-wide Δz^2^ = 0.01 ± 0.02 π^2^). The eight loci overlapped between 2 to 179 genes and together contain 11 candidate genes with known roles in bone, cartilage and/or limb development (e.g., *Nkx3-*2 and *Sox9*; Table 1). Four out of the eight loci contain genes with a “short tibia” or “short limb” knockout phenotype (Table 1; *P* ≤ 0.032 from 1000 permutations, see Methods for details). Of the broader set of genes at these loci with any limb knockout phenotypes, only *fibrillin 2* (*Fbn2*) is polymorphic for SNPs coding for different amino acids, suggesting that for the majority of loci with large shifts in allele frequency, gene regulatory mechanism(s) were likely important in the selection response (fig. S7; Table S3; see Supplementary Notes for further analyses on enrichment in gene functions, protein-coding vs. *cis-*acting changes and clustering with loci affecting human height).

**Fig. 2.**
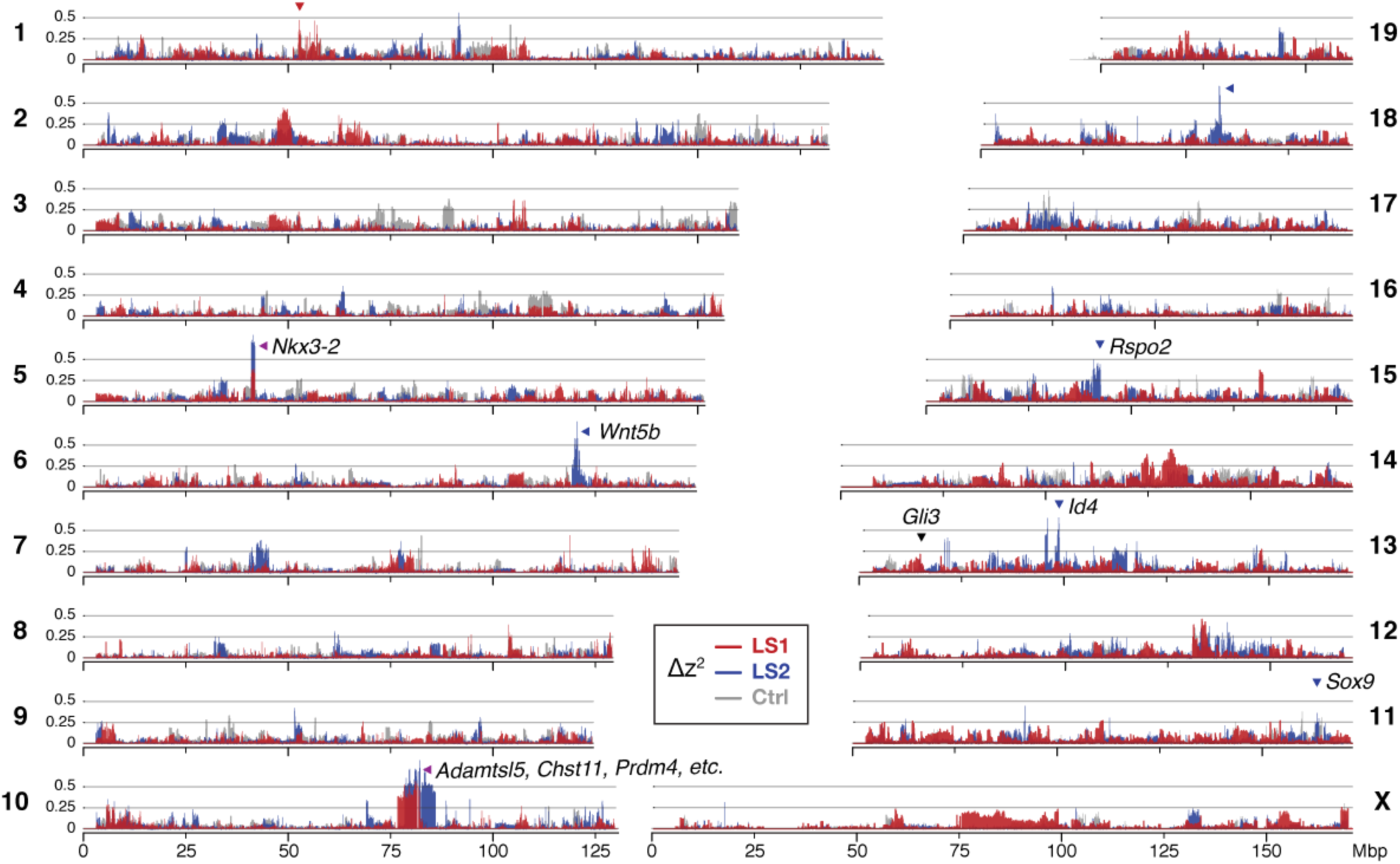
Widespread genomic response to selection for increased tibia length. Allele frequency shifts between generations F0 and F17 in LS1, LS2 and Ctrl lines are shown as Δz^2^ profile across the genome (plotted here as fraction of its range from 0 to π^2^). The Ctrl Δz^2^ profile (grey) confirmed our expectation from theory and simulation that drift, inbreeding and genetic linkage could combine to generate large Δz^2^ shifts even without selection. Nonetheless the LS1 (red) and LS2 (blue) profiles show a greater number of strong and parallel shifts than Ctrl. These selective sweeps provide support for the contribution of discrete loci to selection response (arrowheads, red: LS1; blue: LS2; purple: parallel; see also fig. S4, S5) beyond a polygenic background, which may explain a majority of the selection response but leave little discernible selection signature. Candidate genes are highlighted (Table 1).

Taken together, two major observations stand out from our genomic survey. One, a polygenic, infinitesimal selection model with strong LD amongst marker SNPs best fits the observed data; and two, we nevertheless find more discrete loci in LS1 and LS2 than in Ctrl, beyond the significance threshold set by the infinitesimal model (Fig. 2; fig. S5). Thus, We conclude that the genetic basis of the selection response in the Longshanks experiment has a significant contribution from discrete loci with major effect, but the remaining response may be indistinguishable from a polygenic response.

| Rk | Chr | Span (Mbp) | Peak | | | | $\Delta q$ | | | Type | Candidate genes* |
| --- | --- | --- | --- | --- | --- | --- | --- | --- | --- | --- | --- |
|  |  |  |  | Core (kbp) | TAD (kbp) | Genes | LS1 | LS2 | Ctrl |  |  |
| 1 | 5 | 38.95–45.13 | 41.77 | 900 | 720 | 3 | 0.69 | 0.86 | -0.14 | Parallel | *Nkx3-2* |
| 2 | 10 | 77.47–87.69 | 81.07 | 5,360 | 6,520 | 175 | 0.79 | 0.88 | -0.04 | Parallel | *Sbno5*, *Aes*, *Adamtsl5* *,<br>*Chst11* *, *Cry1*, *Prdm4* * |
| 3 | 18 | 53.63–63.50 | 58.18 | 220 | 520 | 4 | 0.05 | 0.78 | -0.06 | LS2-specific | - |
| 4 | 13 | 35.59–55.21 | 48.65 | 70 | 2,600 | 22 | 0.24 | 0.80 | -0.03 | LS2-specific | *Id4* |
| 5 | 1 | 53.16–57.13 | 55.27 | 10 | 720 | 4 | 0.65 | 0.01 | -0.23 | LS1-specific | - |
| 6 | 15 | 31.92–44.43 | 41.54 | 10 | 680 | 3 | -0.23 | 0.66 | 0.02 | LS2-specific | *Rspo2* * |
| 7 | 6 | 118.65–125.25 | 120.30 | 130 | 1,360 | 12 | -0.03 | 0.79 | -0.15 | LS2-specific | *Wnt5b* * |
| 8 | 11 | 111.10–115.06 | 113.42 | 10 | 2,120 | 2 | -0.14 | 0.66 | -0.15 | LS2-specific | *Sox9* * |
| **Rk** |  | Rank |  |  |  |  |  |  |  |  |  |
| **Chr** |  | Chromosome |  |  |  |  |  |  |  |  |  |
| **Core** | | Span of 10 kbp windows above $H_{INF, \max LD} P \leq 0.05$ significance threshold | | | | | | | | | |

**TAD** Merged span of topologically associating domains (TAD) overlapping the core span. TADs mark segments along a chromosome that share a common regulatory mechanism. Data from (22).
\* Genes within the TAD span showing “short tibia”, “short limbs”, “abnormal osteoblast morphology” or “abnormal cartilage morphology” knockout phenotypes are listed, with \* marking those with “short tibia”.

**Table 1. Major loci likely contributing to the selection response.** These 8 loci show significant allele frequency shifts in Δz^2^ and are ordered according to their estimated selection coefficients. Shown for each locus are the full hitchhiking spans, peak location and their size covering the core windows, the overlapping TAD and the number of genes found in it. The two top-ranked loci show shifts in parallel in both LS 1 & 2, with the remaining six showing line-specific response (LS1: 1; LS2: 5). Candidate genes found within the TAD with limb, cartilage or bone developmental knockout phenotypes functions are shown, with asterisks (*) marking those with a “short tibia” knockout phenotype (see also fig. S6 and Table S3 for full table).

We next tested the repeatability of the selection response at the gene/locus level using the two LS replicates. If the founding populations shared the same selectively favored variants, we may observe parallelism or co-incident selective sweeps, as long as selection could overcome random drift. Indeed, the Δz^2^ profiles of LS 1 & 2 were more similar to each other than to Ctrl (Fig. 2 & 3A; fig. S8; Pearson’s correlation in Δz^2^ from 10 kbp windows: LS1–LS2: 0.21, vs. LS1–Ctrl: 0.063 and LS2–Ctrl: 0.045). Whereas previous genomic studies with multiple natural or artificial selection replicates focused mainly on detecting parallel loci (*23*-*26*), here we have the possibility to quantify parallelism and determine the selection value of a given locus. Six out of eight significant loci at the *H*_*INF, max LD*_ threshold were line-specific, even though the selected alleles were always present in the F0 generation in both lines. This prevalence of line-specific loci was consistent, even if we used different thresholds. However, the two remaining loci that ranked first and second by selection coefficient were parallel, both with *s* > (Fig. 3B; note that as outliers, the selection coefficient may be substantially overestimated, but their rank order should remain the same), supporting the idea that the probability of parallelism can be high among those loci with the greatest selection advantage (*27*). In contrast to changes in global diversity over 17 generations, where we could only detect a slight difference between the LS lines and Ctrl, we found the signature of parallelism to be significantly different between the selected LS1 and LS2 replicates, as opposed to comparisons with Ctrl, or between simulated replicates (fig. S8; *χ* ^2^ test, LS1–LS2: *P* ≤ 1 ×10^-10^; Ctrl–LS1 and Ctrl–LS2, *P* > 0.01 and *P >* 0.2, respectively, both non-significant after correcting for multiple testing; see Supplementary Notes for details). As such, the parallel selected loci between LS1 and LS2 provide the strongest evidence for the role of discrete major loci; and represent prime candidates for molecular dissection (see fig. S9 and Supplementary Notes “Molecular dissection of *Gli3*” for an additional locus).

**Fig. 3.**
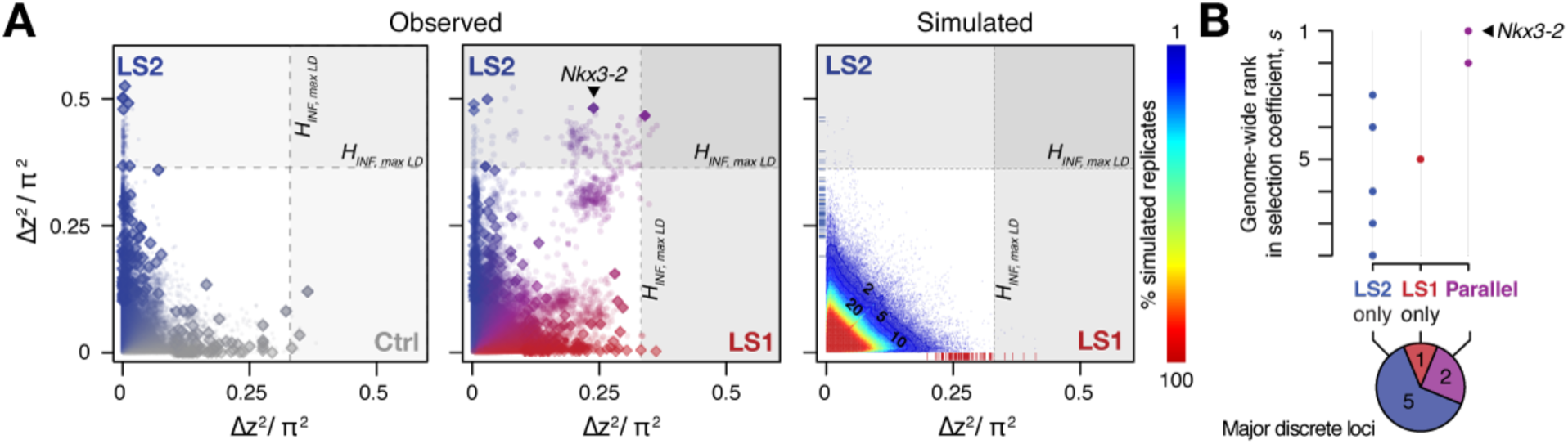
Selection response in the Longshanks lines was largely line-specific, but the strongest signals occurred in parallel. (**A**) Allele frequencies showed greater shifts in LS2 (blue) than in Ctrl (grey; left panel; diamonds: peak windows; dots: other 10 kbp windows; see fig. S8 for Ctrl vs. LS1 and Supplementary Methods for details). Changes in the two lines were not correlated with each other. In contrast, there were many more parallel changes in a comparison between LS1 (red) vs. LS2 (blue; middle panel). The overall distribution closely matches simulated results under the infinitesimal model with maximal linkage disequilibrium (*H*_*INF,* *max* *LD*_; right heatmap summarizes the percentage seen in 100 simulated replicates), with most of the windows showing little to no shift (red hues near 0; see also fig. S8 for an example replicate). Tick marks along axes showed genome-wide maximum Δz^2^ shifts in each replicate in LS1 (red) and LS2 (blue), from which we derived line-specific thresholds at the *P* ≤ 0.05 significance level. While the simulated results showed good match to the bulk of the observed data, it did not recover the strong parallel shifts observed between LS1 and LS2 (compare middle to right panels, points along the diagonal; adjacent windows appear as clusters due to hitchhiking). (**B**) Genome-wide ranking based on estimated selection coefficients *s* among the candidate discrete loci at *P* ≤ 0.05 under *H*_*INF,* *max* *LD*_. While six out of eight total loci showed significant shift in only LS1 or LS2, the two loci top-ranked by selection coefficients were likely selected in parallel in both LS1 and LS2 (also see left panel in A).

### Molecular dissection of the Nkx3-2 locus highlights cis-acting changes

Between the two major parallel loci, we chose the locus on chromosome 5 (Chr5) at 41– 42 Mbp for functional validation because it showed the strongest estimated selection coefficient and both Longshanks lines shared a clear and narrow selection signature. Crucially, it contains only three genes, including *Nkx3-2*, a known regulator of bone maturation (also known as *Bapx1*; Fig. 2 & 4A) (*29*). At this locus, the pattern of variation resembles a selective sweep spanning 1 Mbp (Fig. 5A). Comparison between F0 and F17 individuals revealed no recombinant in this entire region (fig. S11A, top panel), precluding fine-mapping using recombinants. We then analyzed the genes in this region to identify the likely target(s) of selection. First, we determined that no coding changes existed for either *Rab28* or *Nkx3-2,* the two genes located within the topologically associating domain (TADs, which mark chromosome segments with shared gene regulatory logic) (*22*). We then performed *in situ* hybridization and detected robust expression of *Nkx3-2* and *Rab28* in the developing fore-and hindlimb buds of Ctrl, LS1 and LS2 E12.5, in a domain broadly overlapping the presumptive zeugopod, the region including the tibia (fig. S10B). A third gene, *Bod1l*, straddled the TAD boundary with its promoter located in the neighboring TAD, making its regulation by sequences in the selected locus unlikely. Consistent with this, *Bod1l* showed only weak or undetectable expression in the developing limb bud (fig. S10A). We next combined ENCODE chromatin profiles and our own ATAC-Seq data to identify limb enhancers in the focal TAD. Here we found 3 novel enhancer candidates (N1, N2 and N3) carrying 3, 1 and 3 SNPs respectively, all of which showed significant allele frequency shifts in LS1 & 2 (Fig. 4B & C; fig. S11A). Chromosome conformation capture assays showed that the N1–N3 sequences formed long-range looping contacts with the *Nkx3-2* promoter—a hallmark of enhancers—despite nearly 600 kbp of intervening sequence (Fig. 4B). We next used transgenic reporter assays to determine whether these sequences could drive expression in the limbs. Here, we were not only interested in whether the sequence encoded enhancer activity, but specifically whether the SNPs would affect the activity (Fig. 4D). We found that the F0 alleles of the N1 and N3 enhancers (3 SNPs each in about 1 kbp) drove robust and consistent *lacZ* expression in the developing limb buds (N1 and N3) as well as in expanded trunk domains (N3) at E12.5 (Fig. 4E). In contrast, transgenic reporters carrying the selected F17 alleles of N1 and N3 from the Chr 5: 41 Mbp locus showed consistently weak, nearly undetectable *lacZ* expression (Fig. 4E). Thus, switching from the F0 to the F17 enhancer alleles led to a nearly complete loss in activity (“loss-of-function”). This is consistent with the role of *Nkx3-2* as a repressor in long bone maturation (*29*). We hypothesize that the F17 allele causes *de-repression* of bone formation by reducing enhancer activity and *Nkx3-2* expression. Crucially, the F0 N1 enhancer showed activity that presages future long bone cartilage condensation in the limb (Fig. 4D). This pattern recalls previous results that suggest that undetected early expression of *Nkx3-2* may mark the boundaries and size of limb bone precursors, including the tibia (*30*) (fig. S10C). Conversely, over-expression of *Nkx3-2* has been shown to cause shortened tibia (even loss) in mice (*31*, *32*). In humans, homozygous frameshift mutations in *NKX3-2* cause the rare disorder spondylo-megaepiphyseal-metaphyseal dysplasia (SMMD; OMIM: 613330) that is characterized by short-trunk, long-limbed dwarfism and bow-leggedness (*33*). This broadly corresponds to the expression domains of the two novel N1 (limbs) and N3 (limbs and trunk) enhancers. Instead of wholesale loss of *Nkx3-2* expression, which would have been lethal in mice (*34*) or likely cause major defects similar to SMMD patients (*33*), our *in situ* hybridization data did not reveal qualitative differences in *Nkx3-2* expression domains between Ctrl or LS embryos (fig. S10B). Taken together, our results recapitulate the key features of *cis*-acting mode of adaptation: *Nkx3-2* is a broadly expressed pleiotropic transcription factor, which causes lethality when knocked out (*34*). We found no amino acid changes that could impact its protein function. Rather, changes of tissue-specific expression by modular enhancers likely played a more important role. By combining population genetics, functional genomics and developmental genetic techniques, we were able to dissect a megabase-long locus and present data supporting the identification of up to 6 candidate quantitative trait nucleotides (QTNs). In mice, this represents a rare example of genetic dissection of a trait to the base-pair level.

**Fig. 4.**
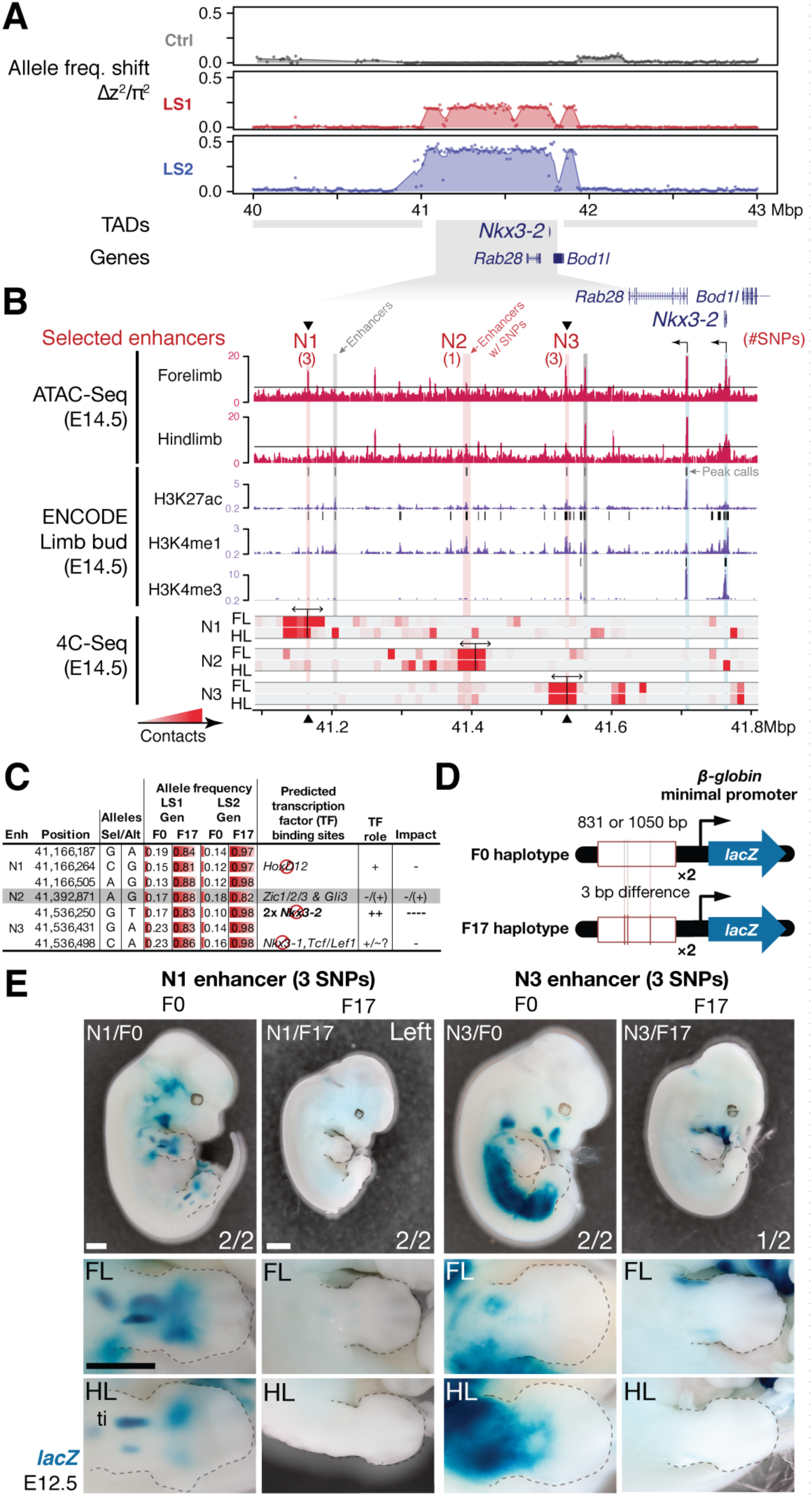
Strong parallel selection response at the bone maturation repressor *Nkx3-2* locus was associated with decreased activity of two enhancers. (**A**) Δz^2^ in this region showed strong parallel differentiation spanning 1 Mbp in both Longshanks but not in the Control line, which contains three genes *Nkx3-2*, *Rab28* and *Bod1l* (whose promoter lies outside the TAD boundary, shown as grey boxes). In LS2, an originally rare allele had almost swept to fixation (fig. S11A & B). (**B**) Chromatin profiles (ATAC-Seq, red, (*28*); ENCODE histone modifications, purple) from E14.5 developing limb buds revealed five putative limb enhancers (grey and red shading) in the TAD, three of which contained SNPs showing significant frequency shifts. Chromosome conformation capture assays (4C-Seq) from E14.5 limb buds from the N1, N2 and N3 enhancer viewpoints (bi-directional arrows) showed significant long-range looping between the enhancers and sequences around the *Nkx3-2* promoter (heat-map with red showing increased contacts; Promoters are shown with black arrows and blue vertical shading). (**C**) SNPs found within the N1–3 enhancers showed about 0.75 increase in allele frequency, with 3 sites leading to gained or lost (red symbols) predicted transcription factor binding sites *Nkx3-2* pathway (including a single site in N3 causing loss of 2 *Nkx3-2* binding site), which are predicted to reduce enhancer activity in N1 and N3. (**D, E**) Transient transgenic reporter assays of the N1 and N3 enhancers showed that the F0 alleles drove robust and consistent expression at centers of future cartilage condensation (N1) and broader domains of *Nkx3-2* expression (N3) in E12.5 fore-and hindlimb buds (FL, HL; ti: tibia). Fractions indicate number of embryos showing similar *lacZ* staining out of all transgenic embryos. Substituting the F17 allele (replacing 3 positions each in N1 and N3) led to little observable limb bud expression in both the N1/F17 and the N3/F17 enhancers, suggesting that selection response for longer tibia involved de-repression of bone maturation through a loss-of-function regulatory allele of *Nkx3-2* at this locus. Scale bar: 1 mm for both magnifications.

**Figure 5.**
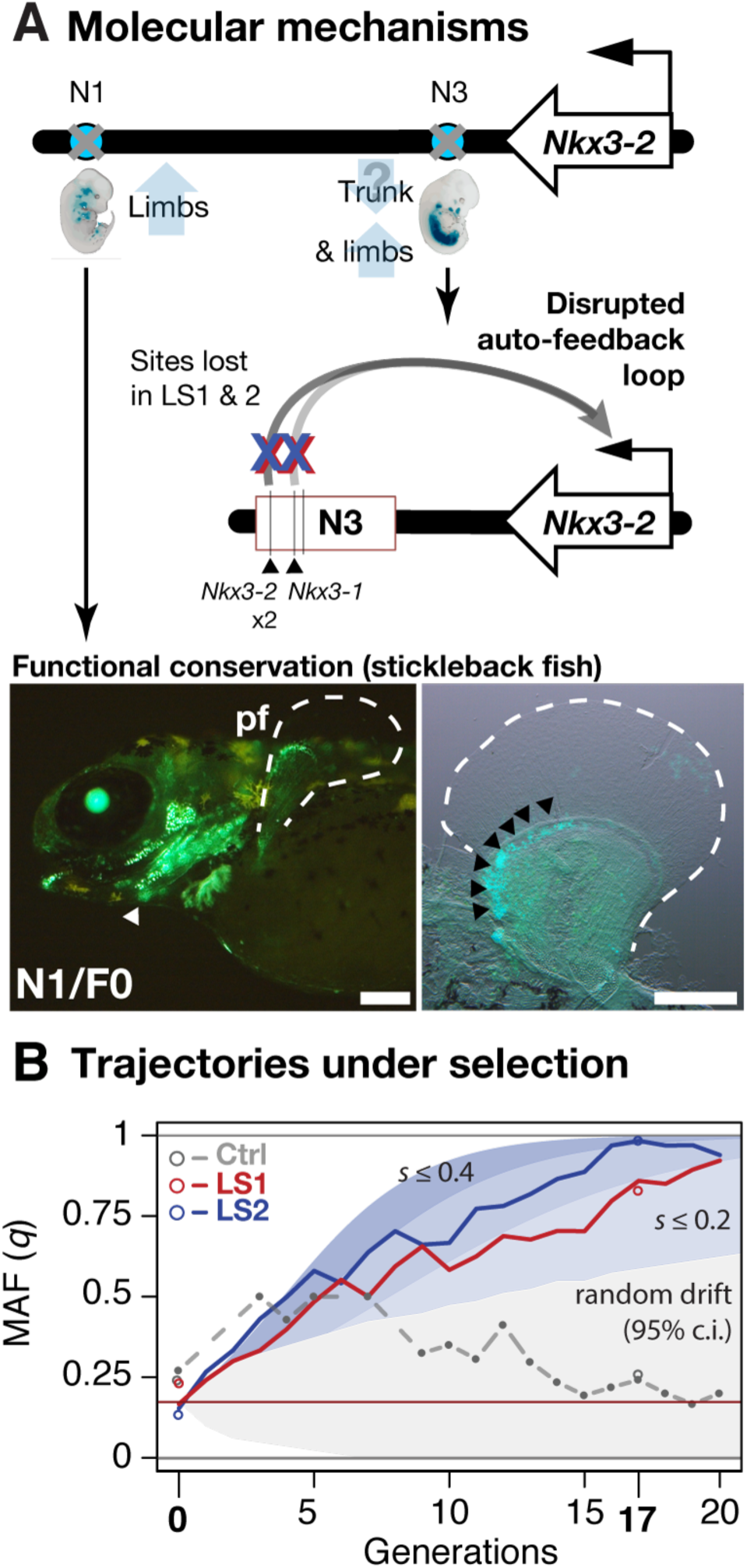
Linking base-pair changes to rapid morphological evolution. (**A**) At the *Nkx3-2* locus, we identified two long-range enhancers (circles), N1 and N3, located 600 and 230 kbp away, respectively. They drive partially overlapping expression domains in limbs (N1 and N3) and trunk (N3) during development, sites that may correlate positively (tibia length) and possibly negatively (trunk with body mass) with the Longshanks selection regime. For both enhancers, the selected F17 alleles carry loss-of-function variants (grey crosses). Two out of three SNPs in the N3 enhancer are predicted to disrupt an auto-feedback loop, likely reducing *Nkx3-2* expression in the trunk and limb regions. Similarly, the enhancer function of the strong F0 allele evolutionarily conserved in fishes, demonstrated by its ability to drive consistent GFP expression (green) in the pectoral fins (pf, outlined) and branchial arches (arrowhead) in transgenic stickleback embryos at 11 days post-fertilization. The N1 enhancer can recapitulate *nkx3.2* expression in distal cells in the endochondral radial domain in developing fins (black arrowheads, right). Scale bar: 250 µm for both magnifications. (**B**) Allele frequency of the selected allele (minor F0 allele, *q*) at N3 over 20 generations (red: LS1; blue: LS2; grey broken line: Ctrl; results from N1 were nearly identical due to tight linkage). Observed frequencies from genotyped samples in the Ctrl line are marked with filled circles. Dashed lines indicate missing Ctrl generations. Open circles at generations 0 and 17 indicate allele frequencies from whole genome sequencing. The allele frequency fluctuated in Ctrl due to random drift, whereas there was a generally linear increase from around 0.17 to 0.85 (LS1) and 0.98 (LS2) by generation 17. Shaded contours mark trajectories under varying selection coefficients starting from 0.17 over time (the average between LS1 & LS2 founders). The grey shaded region mark the 95% confidence interval under random drift.

### Linking molecular mechanisms to evolutionary consequence

We next aimed to determine the evolutionary relevance of the *Nkx3-2* enhancer variants at the molecular and the population level. At the strongly expressed N3/F0 “trunk and limb” enhancer, we note that the SNPs in the F17 selected allele lead to disrupted *Nkx3-1* and *Nkx3-2* binding sites [Fig. 4C & 5A; UNIPROBE database (*35*)]. This suggests that the selected SNPs may disrupt an auto-feedback loop to decrease *Nkx3-2* activity in the limb bud and trunk domain (Fig. 5A). Using a *GFP* transgenic reporter assay in stickleback fish embryos, we found that the mouse N1/F0 enhancer allele was capable of driving expression in the distal cells but not in the fin rays of the developing fins (Fig. 5A). This pattern recapitulates fin expression of *nkx3.2* in fish, which gives rise to endochondral radials (homologous to ulna/tibia in mice) (*36*). Our results suggest that despite its deep functional conservation, strong selection may have favored the weaker N1/F17 and N3/F17 enhancer alleles in the context of the Longshanks selection regime.

Using theory and simulations, we can go beyond the qualitative level and quantitatively estimate the selection coefficient and the contribution of the *Nkx3-2* locus to the total selection response in the Longshanks mice. We retraced the selective sweep of the *Nkx3-2* N1 and N3 alleles through targeted genotyping in 1569 mice across all 20 generations. The selected allele steadily increased from around 0.17 to 0.85 in LS1 and 0.98 in LS2 but fluctuated around 0.25 in Ctrl (Fig. 5B). We estimate that such a change of around 0.8 in allele frequency would correspond to a selection coefficient *s* of ∼0.24 ± 0.12 at this locus (Fig. 5C; see Supplementary Notes section on “Estimating selection coefficient”). By extending our simulation framework to allow for a major locus against an infinitesimal background, we find that the *Nkx3-2* locus would have to contribute 9.4% of the total selection response (limits 3.6 – 15.5%; see Supplementary Notes section “Estimating selection coefficient” for details) in order to produce a shift of 0.8 in allele frequency over 17 generations.

## Discussion

A defining task of our time is to understand the factors that determine and constrain how small populations respond to sudden environmental changes. Here, we analyze the genomic changes in the Longshanks experiment, which was conducted under replicated and controlled conditions, to characterize the genomic changes that occur as small experimental populations respond to selection.

An important conclusion from the Longshanks experiment is that tibia length increased readily and repeatedly in response to selection even in an extremely bottlenecked population with as few as 14-16 breeding pairs. This is because the lines were founded with enough standing variation, and generation 17 was still only a fraction of the way to the characteristic time for the selection response at ∼2*N*_*e*_ generations (*37*), estimated here to be around 90 (fig. S2B; see Supplementary Notes on simulation). This *N*_*e*_ of 46, while small, is comparable to those in natural populations like the Soay sheep (35.3), Darwin’s finches (38–60) or Tasmanian Devils (26–37; this last study documents a rapid and parallel evolutionary response to transmissible tumor) (*38*-*40*). Our results underscore the importance of standing genetic variation in rapid adaptive response to a changing environment, a recurrent theme in natural adaptation (*24*, *39*, *41*) and breeding (*42*).

By combining pedigree records with sequencing of founder individuals, our data had sufficient detail to allow precise modeling of trait response with predicted shifts in allele frequency distribution that closely matched our results, and with specific loci that we functionally validated. Our results imply a mixed genetic architecture with a few discrete loci of large effect, amid an infinitesimal background. This finding highlights another advantage of evolve-and-resequence (E&R) experiments over quantitative trait loci (QTL) mapping crosses: by sampling a much broader pool of alleles and continually competing them against each other, the inferred genetic architecture and distribution of effect sizes are more likely to be representative of the population at large.

Parallel evolution is often seen as a hallmark for detecting selection (*25*, *43*-*45*). We investigated the factors favoring parallelism by contrasting the two Longshanks replicates against the Control line. We observed little to no parallelism between selected lines and Ctrl, or between simulated replicates of selection, even though the simulated haplotypes were sampled directly from actual founders. This underscores that parallelism depends on both shared selection pressure (absent in Ctrl) *and* the availability of large effect loci that confer a substantial selection advantage (absent under the infinitesimal model; Fig. 3A&B; fig. S8).

Through in-depth dissection of the *Nkx3-2* locus, our data show in fine detail how the selective value of standing variants depends strongly on the selection regime: the originally common F0 variant of the N1 enhancer show deep functional conservation and can evidently recapitulate fin *nkx3.2* expression in fishes (Fig. 5A). Yet in the Longshanks experiment selection strongly favored the inactive allele (Fig. 5B). Similarly, our molecular dissection of two loci show that both gain-of-function (*Gli3*) and loss-of-function (*Nkx3-2*) variants could be favored by selection (Fig. 4E, 5A; fig. S9D). Through synthesis of multiple lines of evidence, our work uncovered the key role of *Nkx3-2*, which was not an obvious candidate gene like *Gli3* due to the lack of limb phenotype in the *Nkx3-2* knockout mice. To our surprise, the same loss of *NKX3-2* function in human SMMD patients manifests itself in opposite ways in different bone types as short trunk and long limbs (*33*). This matches the expression domains of our N1 (limb) and N3 (limb and trunk) enhancers (Fig. 5A). In the absence of any lethal coding mutations, evidently the F17 haplotype carried beneficial alleles at both enhancers for the limb and potentially also trunk target tissues; and was therefore strongly favored under the novel selection regime in the Longshanks selection experiment. We estimate that these enhancer variants, along with any other tightly linked beneficial SNPs, segregate as a single locus and contributes ∼10% of the overall selection response.

## Conclusion

Using the Longshanks selection experiment, we show that by synthesizing theory, empirical data and molecular genetics, it is possible to identify some of the individual SNPs that have contributed to the response to selection on morphology. In particular, discrete, large effect loci are revealed by their parallel response. Further work should focus on dissecting the mechanisms behind the dynamics of selective sweeps and/or polygenic adaptation by resequencing the entire selection pedigree; testing how the selection response depends on the genetic architecture; and the extent to which linkage places a fundamental limit on our inference of selection. Improved understanding in these areas may have broad implications for conservation, rapid adaptation to climate change and quantitative genetics in medicine, agriculture and in nature.

## Acknowledgements

We thank Felicity Jones for input into experimental design, helpful discussion and for improving the manuscript. We thank the Rolian, Barton, Chan and Jones Labs members for support, insightful scientific discussion and improving the manuscript. We thank the Rolian lab members, the Animal Resource Centre staff at the University of Calgary and MPI Dresden Animal Facility staff for animal husbandry. We thank Derek Lundberg for help with library preparation automation. We thank Christa Lanz, Rebecca Schwab and Ilja Bezrukov for assistance with high-throughput sequencing and associated data processing; Andre Noll for high-performance computing support; the MPI Tübingen IT team for computational support. We thank Felicity Jones and the Jones Lab for help with stickleback microinjections. pRS16 was a gift from François Spitz. We thank Mirna Marinič for creating an earlier version of the transgenic reporter plasmid. We are indebted to Gemma Puixeu Sala, William G. Hill, Peter Keightley for input and discussion on data analysis and simulation. We are also indebted to Stefan Mundlos, Przemko Tylzanowki, Weikuan Gu for suggested experiments and sharing unpublished data. We thank Sean B. Carroll, Andrew Clark, Jonathan Pritchard, Matthew Rockman, Gregory Wray and David Kingsley for thoughtful input that has greatly improved our manuscript. J.P.L.C. is supported by the International Max Planck Research School “From Molecules to Organisms”. S.B. and N.B. are supported by IST Austria. C.R. is supported by Discovery Grant #4181932 from the Natural Sciences and Engineering Research Council of Canada and by the Faculty of Veterinary Medicine at the University of Calgary. Y.F.C. is supported by the Max Planck Society.

## Author Contributions

C.R. designed and initiated the Longshanks selection experiment. M.M. and C.R. performed the selection, phenotyping and collected tissue samples for sequencing. Y.F.C. and C.R. designed the sequencing strategy. W.H.B., M.K., prepared the samples and performed sequencing. S.B., N.B. performed simulations and analyzed data. M.M., S.B., N.B., C.R., Y.F.C. analyzed the pedigree data. J.P.L.C., M.N.Y., M.K., I.S., J.C., C.R. and Y.F.C. designed, performed and analyzed results from functional experiments. J.P.L.C., R.N. and Y.F.C. planned and performed and analyzed the mouse transgenic experiments. S.B., M.N.Y., C.R., N.B. and Y.F.C analyzed the genomic data. All authors discussed the results and implications, wrote and commented on the manuscript at all stages.

## Materials and Methods

### Animal Care and Use

All experimental procedures described in this study have been approved by the applicable University institutional ethics committee for animal welfare at the University of Calgary (HSACC Protocols M08146 and AC13-0077); or local competent authority: Landesdirektion Sachsen, Germany, permit number 24-9168.11-9/2012-5.

### Reference genome assembly

All co-ordinates in the mouse genome refer to *Mus musculus* reference mm10, which is derived from GRCm38.

### Code and data availability

Sequence data have been deposited in the GEO database under accession number `[X]`. Non-sequence data have been deposited at Dryad under accession number `[Y]`. Analytical code and additional notes have been deposited in the following repository: https://github.com/evolgenomics/Longshanks.

### Pedigree data

Tibia length and body weight phenotypes were measured as previously described (*14*). A total of 1332 Control, 3054 LS1, and 3101 LS2 individuals were recorded. Five outlier individuals with a skeletal dysplasia of unknown etiology were removed from LS2 and excluded from further analysis. Missing data in LS2 were filled in with random individuals that best matched the pedigree. Trait data were analyzed to determine response to selection based on the measured traits and their rank orders based on the selection index.

### Simulations

Simulations were based on the actual pedigree and selection scheme, following one chromosome at a time. Each chromosome was represented by a set of junctions, which recorded the boundaries between genomes originating from different founder genomes; at the end, the SNP genotype was reconstructed by seeding each block of genome with the appropriate ancestral haplotype. This procedure is much more efficient than following each of the very large number of SNP markers. Crossovers were uniformly distributed, at a rate equal to the map length (*46*). Trait value was determined by a component due to an infinitesimal background (*V*_*g*_); a component determined by the sum of effects of 10^4^ evenly spaced discrete loci (*V*_*s*_); and a Gaussian non-genetic component (*V*_*e*_). The two genetic components had variance proportional to the corresponding map length, and the heritability was estimated from the observed trait values (see Supplementary Notes under “Simulations”). In each generation, the actual number of male and female offspring were generated from each breeding pair, and the male and female with the largest trait value were chosen to breed.

SNP genotypes were assigned to the founder genomes with their observed frequencies. However, to reproduce the correct variability requires that we assign founder *haplotypes*. This is not straightforward, because low-coverage individual genotypes cannot be phased reliably, and heterozygotes are frequently mis-called as homozygotes. We compared three procedures, which were applied within intervals that share the same ancestry: assigning haplotypes in linkage equilibrium (LE, or “no LD”); assigning heterozygotes to one or other genome at random, which minimises linkage disequilibrium, given the diploid genotype (“min LD”); and assigning heterozygotes consistently within an interval, which maximises linkage disequilibrium (“max LD”) (fig. S2C). For details, see Supplementary Notes.

### Significance thresholds

To obtain significance thresholds, we summarize the genome-wide maximum Δz^2^ shift for each replicate of the simulated LS1 and LS2 lines, averaged within 10kb windows, and grouped by the selection intensity and extent of linkage disequilibrium (LD). From this distribution of genome-wide maximum Δz^2^ we obtained the critical value for the corresponding significance threshold (typically the 95^th^ quantile or *P* = 0.05) under each selection and LD model (Fig. 3A; fig. S2E). This procedure controls for the effect of linkage and hitchhiking, line-specific pedigree structure, and selection strength.

### Sequencing, genotyping and phasing pipeline

Sequencing libraries for high-throughput sequencing were generated using TruSeq or Nextera DNA Library Prep Kit (Illumina, Inc., San Diego, USA) according to manufacturer’s recommendations or using equivalent *Tn5* transposase expressed in-house as previously described (*47*). Briefly, genomic DNA was extracted from ear clips by standard Protease K digestion (New England Biolabs GmbH, Frankfurt am Main, Germany) followed by AmpureXP bead (Beckman Coulter GmbH, Krefeld, Germany) purification. Extracted high-molecular weight DNA was sheared with a Covaris S2 (Woburn, MA, USA) or “tagmented” by commercial or purified *Tn5*-transposase according to manufacturer’s recommendations. Each sample was individually barcoded (single-indexed as N501 with N7XX variable barcodes; all oligonucleotides used in this study were synthesized by Integrated DNA Technologies, Coralville, Iowa, USA) and pooled for high-throughput sequencing by a HiSeq 3000 (Illumina) at the Genome Core Facility at the MPI Tübingen Campus. Sequenced data were pre-processed using a pipeline consisting of data clean-up, mapping, base-calling and analysis based upon fastQC v0.10.1 (*52*); trimmomatic v0.33 (*48*); bwa v0.7.10-r789 (*49*); GATK v3.4-0-gf196186 modules BQSR, MarkDuplicates, IndelRealignment (*50*, *51*). Genotype calls were performed using the GATK HaplotypeCaller under the GENOTYPE_GIVEN_ALLELES mode using a set of high-quality SNP calls made available by the Wellcome Trust Sanger Centre (Mouse Genomes Project version 3 dbSNP v137 release (*52*), after filtering for sites segregating among inbred lines that may have contributed to the original 7 female and 2 male CD-1 founders, namely 129S1/SvImJ, AKR/J, BALB/cJ, BTBR *T*^*+*^ *Itpr3*^*tf*^/J, C3H/HeJ, C57BL/6NJ, CAST/EiJ, DBA/2J, FVB/NJ, KK/HiJ, MOLF/EiJ, NOD/ShiLtJ, NZO/HlLtJ, NZW/LacJ, PWK/PhJ and WSB/EiJ based on (*17*). We consider a combined ∼100x coverage sufficient to recover any of the 18 CD-1 founding haplotypes still segregating at a given locus. The raw genotypes were phased with Beagle v4.1 (*53*) based on genotype posterior likelihoods using a genetic map interpolated from the mouse reference map (*46*) and imputed from the same putative CD-1 source lines as the reference panel. The site frequency spectra (SFS) were evaluated to ensure genotype quality (fig. S3A).

### Population genetics summary statistics

Summary statistics of the F0 and F17 samples were calculated genome-wide [Weir–Cockerham F_ST_, π, heterozygosity]; in adjacent 10 kbp windows (Weir–Cockerham F_ST_, π, allele frequencies *p* and *q*), or on a per site basis (Weir–Cockerham F_ST_, π, *p* and *q*) using `VCFtools v0.1.14` (*54*). The summary statistics Δz^2^ was the squared within-line difference in arcsine square root transformed MAF *q*; it ranges from 0 to π^2^. The resulting data were further processed by custom bash, Perl and R v3.2.0 (*55*) scripts.

### Peak loci and filtering for hitchhiking windows

Peak loci were defined by a descending rank ordering of all 10 kbp windows, and from each peak signals the windows were extended by 100 SNPs to each side, until no single SNP rising above a Δz^2^ shift of 0.2 π^2^ was detected. A total of 810 peaks were found with a Δz^2^ shift ≥ 0.2 for LS1 & 2. Following the same procedure, we found 766 peaks in Ctrl.

### Candidate genes

To determine whether genes with related developmental roles were associated with the selected variants, the topologically associating domains (TADs) derived from mouse embryonic stem cells as defined elsewhere (*22*) were re-mapped onto mm10 co-ordinates. Genes within the TAD overlapping within 500 kbp of the peak window (“core span”) were then cross-referenced against annotated knockout phenotypes (Mouse Genome Informatics, http://www.informatics.jax.org). This broader overlap was chosen to account for genes whose regulatory sequences like enhancers but not their gene bodies fall close to the peak window. We highlight candidate genes showing limb-and bone-related phenotypes, e.g., with altered limb bone lengths or epiphyseal growth plate morphology, as observed in Longshanks mice (*16*), of the following categories (along with their Mammalian Phenotype Ontology term and the number of genes): “abnormal tibia morphology/MP:0000558” (212 genes), “short limbs/MP:0000547” and “short tibia/MP:0002764” (223 genes), “abnormal cartilage morphology/MP:0000163” (321 genes), “abnormal osteoblast morphology/MP:0004986” (122 genes). Note that we exclude compound mutants or those conditional mutant phenotypes involving transgenes. To determine if the overlap with these genes are significant, we performed 1000 permutations of the core span using `bedtools v2.22.1 shuffle with the -noOverlapping` option (*56*) and excluding ChrY, ChrM and the unassembled scaffolds. We then followed the exact procedure as above to determine the number of genes in the overlapping TAD belonging to each category. We reported the quantile rank as the *P*-value, ignoring ties. To determine other genes in the region, we list all genes falling within the entire hitchhiking window (Table S3).

### Identification of putative limb enhancers

We downloaded publicly available chromatin profiles, derived from E14.5 limbs, for the histone H3 lysine-4 (K4) or lysine-27 (K27) mono-/tri-methylation or acetylation marks (H3K4me1, H3K4me3 and H3K27ac) generated by the ENCODE Consortium (*57*). We intersected the peak calls for the enhancer-associated marks H3K4me1 and H3K27ac and filtered out those overlapping promoters [H3K4me3 and promoter annotation according to the FANTOM5 Consortium (*58*)].

### Enrichment analysis

To calculate enrichment through the whole range of Δz^2^, a similar procedure was taken as in *Candidate genes* above. For knockout gene functions, genes contained in TADs within 500 kbp of peak windows were included in the analysis. We use the complete database of annotated knockout phenotypes for genes or spontaneous mutations, after removing phenotypes reported under conditional or polygenic mutants. For gene expression data, we retained all genes which have been reported as being expressed in any of the limb structures, by tracing each anatomy ontological term through its parent terms, up to the top level groupings, e.g., “limb”, in the Mouse Genomic Informatics Gene Expression Database (*59*). For E14.5 enhancers, we used a raw 500 kbp overlap with the peak windows, because enhancers, unlike genes, may not have intermediaries and may instead represent direct selection targets.

For coding mutations, we first annotated all SNPs for their putative effects using `snpEff v4.0e` (*60*). To accurately capture the per-site impact of coding mutations, we used per-site Δz^2^ instead of the averaged 10 kbp window. For each population, we divided all segregating SNPs into up to 0.02 bands based on per-site Δz^2^. We then tracked the impact of coding mutations in genes *known to be expressed in limbs*, as above. We reported the sum of all missense (“moderate” impact), frame-shift, stop codon gain or loss sites (“high impact”). A linear regression was used to evaluate the relationship between Δz^2^ and the average impact of coding SNPs (SNPs with high or moderate impact to all coding SNPs).

For regulatory mutations, we used the same bins spanning the range of Δz^2^, but focused on the subset of SNPs falling within the ENCODE E14.5 limb enhancers. We then obtained a weighted average conservation score based on an averaged phastCons (*61*) or phyloP (*62*) score in ±250 bp flanking the SNP, calculated from a 60-way alignment between placental mammal genomes [downloaded from the UCSC Genome Browser (*63*)]. We reported the average conservation score of all SNPs within the bin and fitted a linear regression on log-scale. In particular, phastCons scores range from 0 (un-conserved) to 1 (fully conserved), whereas phyloP is the |*log*_10_| of the *P-*value of the phylogenetic tree, expressed as a positive score for conservation and a negative score for lineage-specific accelerated change. We favored using phastCons for its simpler interpretation.

### Impact of coding variants

Using the same SNP effect annotations described in the section above, we checked whether any specific SNP with significant site-wise Δz^2^ in either LS1 or LS2 cause amino acid changes or protein disruptions and are known to cause limb defects when knocked out. For each position we examined outgroup sequences using the 60-way placental mammal alignment to determine the ancestral amino acid state and whether the selected variant was consistent with purifying vs. diversifying selection. The resulting 12 genes that match these criteria are listed in Table S4.

### Association with human height loci

To test if loci known to be associated with human height are clustered with the selected loci in the LS lines, we downloaded the set of previously published 697 SNPs (*64*). In order to facilitate mapping to mouse co-ordinates, each SNP was expanded to 100 kbp centering on the SNP and converted to mm10 positions using the `liftOver tool` with the multiple mapping option disabled (*63*). We were able to assign positions in 655 out of the 697 total SNPs. Then for each of the 810 loci above the *H*_*INF,* *no* *LD*_ threshold the minimal distance to any of the mapped human loci was determined using `bedtools closest` with the −d option (*56*). Should a region actually overlap, a distance of 0 bp was assigned. To generate the permuted set, the 810 loci were randomly shuffled across the mouse autosomes using the bedtools shuffle program with the - `noOverlapping` option. Then the exact same procedure as the actual data was followed to determine the closest interval. The resulting permuted intervals follow an approximately normal distribution, with the actual observed results falling completely below the range of permuted results.

### *In situ* hybridization

Detection of specific gene transcripts were performed as previously described in (*65*). Probes against *Nkx3-2*, *Rab28*, *Bod1l* and *Gli3* were amplified from cDNA from wildtype C57BL/6NJ mouse embryos (Table S5). Amplified fragments were cloned into pJET1.2/blunt plasmid backbones in both sense and anti-sense orientations using the CloneJET PCR Kit (Thermo Fisher Scientific, Schwerte, Germany) and confirmed by Sanger sequencing using the included forward and reverse primers. Probe plasmids have also been deposited with Addgene. *In vitro* transcription from the T7 promoter was performed using the MAXIscript T7 *in vitro* Transcription Kit (Thermo Fisher Scientific) supplemented with Digoxigenin-11-UTP (Sigma-Aldrich) (MPI Tübingen), or with T7 RNA polymerase (Promega) in the presence of DIG RNA labelling mix (Roche) (University of Calgary). Following TURBO DNase (Thermo Fisher Scientific) digestion probes were cleaned using SigmaSpin Sequencing Reaction Clean-Up columns (Sigma-Aldrich) (MPI Tübingen), or using Illustra MicroSpin G-50 columns (GE Healthcare) (University of Calgary). During testing of probe designs, sense controls were used in parallel reactions to establish background non-specific binding.

### ATAC-seq library preparation and sequencing pipeline

ATAC-seq was performed on dissected C57BL/6NJ E14.5 forelimb and hindlimb. Nuclei preparation and tagmentation were performed as previously described in (*28*), with the following modifications. To minimize endogenous protease activity, cells were strictly limited to 5 + 5 minutes of collagenase A treatment at 37 °C, with frequent pipetting to aid dissociation into single-cell suspensions. Following wash steps and cell lysis, 50 000 nuclei were tagmented with expressed *Tn5* transposase. Each tagmented sample was then purified by MinElute columns (Qiagen) and amplified with Q5 High-Fidelity DNA Polymerase (New England Biolabs) using a uniquely barcoded i7-index primer (N701-N7XX) and the N501 i5-index primer. PCR thermocycler programs were 72°C for 4 min, 98°C for 30 s, 6 cycles of 98°C for 10 s, 65°C for 30 s, 72°C for 1 min, and final extension at 72°C for 4 min. PCR-enriched samples were taken through a double size selection with PEG-based SPRI beads (Beckman Coulter) first with 0.5X ratio of PEG/beads to remove DNA fragments longer than 600 bp, followed by 1.8X PEG/beads ratio in order to select for Fraction A as described in (*66*). Pooled libraries were run on the HiSeq 3000 (Illumina) at the Genome Core Facility at the MPI Tübingen Campus to obtain 150 bp paired end reads, which were aligned to mouse mm10 genome using `bowtie2 v.2.1.0` (*67*). Peaks were called using `MACS14 v.2.1` (*68*).

### Multiplexed chromosome conformation capture (4C-Seq)

Chromosome conformation capture (3C) template was prepared from pooled E14.5 liver, forelimb and hindlimb buds (n = 5–6 C57BL/6NJ embryos per replicate), with improvements to the primer extension and library amplification steps following (*69*). The template was amplified with Q5 High-Fidelity Polymerase (New England Biolabs GmbH, Frankfurt am Main, Germany) using a 4C adapter-specific primer and a pool of 6 *Nkx3-2* enhancer viewpoint primers [and, in a separate experiment, a pool of 8 *Gli3* enhancer-specific viewpoint primers; Table S6]. Amplified fragments were prepared for Illumina sequencing by ligation of TruSeq adapters, followed by PCR enrichment. Pooled libraries were sequenced by a HiSeq 3000 (Illumina) at the Genome Core Facility at the MPI Tübingen Campus with single-end, 150 bp reads. Sequence data were processed using a pipeline consisting of data clean-up, mapping, and analysis based upon cutadapt v1.10 (*70*); bwa v0.7.10-r789 (*49*) ; samtools v1.2 (*71*); bedtools (*56*) and R v3.2.0 (*55*). Alignments were filtered for ENCODE blacklisted regions (*72*) and those with MAPQ scores below 30 were excluded from analysis. Filtered alignments were binned into genome-wide *BglII* fragments, normalized to Reads Per Kilobase of transcript per Million mapped reads (RPKM), and plotted and visualized in R.

### Plasmid construction

Putative limb enhancers corresponding to the F0 and F17 alleles of the *Gli3* G2 and *Nkx3-2* N1 and N3 enhancers were amplified from genomic DNA of Longshanks mice from the LS1 F0 (9 mice) and F17 (10 mice) generations and sub-cloned into pJET1.2/blunt plasmid backbone using the CloneJET PCR Kit (Thermo Fisher Scientific) and alleles were confirmed by Sanger sequencing using the included forward and reverse primers (Table S7). Each allele of each enhancer was then cloned as tandem duplicates with junction *SalI* and *XhoI* sites upstream of a β-globin minimal promoter in our reporter vector (see below). Constructs were screened for the enhancer variant using Sanger sequencing. All SNPs were further confirmed against the rest of the population through direct amplicon sequencing.

The base reporter construct pBeta-lacZ-attBx2 consists of a β-globin minimal promoter followed by a *lacZ* reporter gene derived from pRS16, with the entire reporter cassette flanked by double *attB* sites. The pBeta-lacZ-attBx2 plasmid and its full sequence have been deposited and is available at Addgene.

### Pronuclear injection of F0 and F17 enhancer-reporter constructs in mice

The reporter constructs containing the appropriate allele of each of the 3 enhancers were linearized with *ScaI* (or *BsaI* in the case of the N3 F0 allele due to the gain of a *ScaI* site) and purified. Microinjection into mouse zygotes was performed essentially as described (*73*). At 12 d after the embryo transfer, the gestation was terminated and embryos were individually dissected, fixed in 4% paraformaldehyde for 45 min and stored in PBS. All manipulations were performed by R.N. or under R.N.’s supervision at the Transgenic Core Facility at the Max Planck Institute of Cell Biology and Genetics, Dresden, Germany. Yolk sacs from embryos were separately collected for genotyping and all embryos were stained for *lacZ* expression as previously described (*74*). Embryos were scored for *lacZ* staining, with positive expression assigned if the pattern was consistently observed in at least two embryos.

### *Genotyping of time series at the* Nkx3-2 *N3* locus

Allele-specific primers terminating on SNPs that discriminate between the F0 from the F17 N3 enhancer alleles were designed (rs33219710 and rs33600994; Table S8). The amplicons were optimized as a qPCR reaction to give allele-specific, present/absent amplifications (typically no amplification for the absent allele, otherwise average ΔCt > 10). Genotyping on the entire breeding pedigree of LS1 (n = 602), LS2 (n = 579) and Ctrl (n = 389) was performed in duplicates for each allele on a Bio-Rad CFX384 Touch instrument (Bio-Rad Laboratories GmbH, Munich, Germany) with SYBR Select Master Mix for CFX (Thermo Fisher Scientific) and the following qPCR program: 50°C for 2 min, 95°C for 2 min, 40 cycles of 95°C for 15 s, 58°C for 10 s, 72°C for 10 s. In each qPCR run we included individuals of each genotype (LS F17 selected homozygotes, heterozygotes and F0 major allele homozygotes). For the few samples with discordant results between replicates, DNA was re-extracted and re-genotyped or otherwise excluded.

### Transgenic reporter assays in stickleback fish

In sticklebacks, transgenic reporter assays were carried out using the reporter construct pBHR (*44*). The reporter consists of a zebrafish *heat shock protein 70* (*Hsp70*) promoter followed by an *eGFP* reporter gene, with the entire reporter cassette flanked by *tol2* transposon sequences for transposase-directed genomic integration. The *Nkx3-2* N1/F0 enhancer allele was cloned as tandem duplicates using the *NheI* and *EcoRV* restriction sites upstream of the *Hsp70* promoter. Enhancer orientation and sequence was confirmed by Sanger sequencing. Transient transgenic stickleback embryos were generated by co-microinjecting the plasmid (final concentration: 10 ng/µl) and *tol2* transposase mRNA (40 ng/µl) into freshly fertilized eggs at the one-cell stage as described in (*44*).

## Supplementary Methods

### Supplementary Notes

#### Major considerations in constructing the simulations

In the Longshanks experiment, the highest-ranking male and the highest-ranking female from each family were chosen to breed with the highest-ranking mice from other families within a line (i.e., disallowing sibling matings). Thus, if we disregard non-Mendelian segregation, and the fraction of failed litters (15%), selection acts solely within families, on the measured traits. Such selection does not distort the pedigree, and allows us to follow the evolution of each chromosome separately.

Our simulations track the inheritance of continuous genomes by following the junctions between regions with different ancestry. In principle, we should simulate selection under the infinitesimal model by following the contributions to the trait of continuous blocks of chromosomes across the whole genome. However, this is computationally challenging, since the contributions of all the blocks defined by every recombination event have to be tracked. Instead, we follow a large number of discrete biallelic loci checking that the number is sufficiently large to approach the infinitesimal limit (fig. S2D). We made a further slight approximation by only explicitly modelling discrete loci on one chromosome at a time. We divided the breeding value of an individual into two components. The first, *V*_*g*_, is a contribution from a large number of unlinked loci, due to genes on all but the focal chromosome, as represented by the infinitesimal model. The values of this component amongst offspring are normally distributed around the mean of the parents, with its variance being:

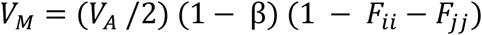

where: *V*_*A*_ is the initial genetic variance, and

*F*_*ii*_,*F*_*jj*_ are the probabilities of identity between distinct genes in each parent,*i*, *j*;

*F*_*ii*_,*F*_*jj*_ are calculated from the pedigree;

β is the fraction of genome on the focal chromosome.

The second component, *V*_*s*_, is the sum of contributions from a large number, *n*, of discrete loci, evenly spaced along the focal chromosome (here we used 10,000), and contributing a fraction *β* of the initial additive variance. We choose these to have equal effects and random signs, ±α, such that initial allele frequencies 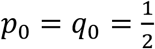, and equal effects α, such that 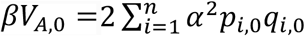. The initial population consists of 28 diploid individuals, matching the experiment, and loci have initial frequencies of 1, 4, 12 and 28 out of the diploid total of 56 alleles, in equal proportions. Inheritance is assumed to be autosomal, with no sex-linkage. This choice of equal effects approaches most closely to the infinitesimal model, for a given number of loci.

The decrease in genetic variance due to random drift is measured by the inbreeding coefficient, defined as the probability of identity by descent, relative to the initial population. We distinguish the identity between two distinct genes within a diploid individual, *F*_*w*_,from the probability of identity between two genes in different individuals, *F*_b_. The overall mean identity between two genes chosen independently and at random from all 2*N* genes is 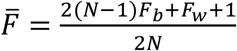. The proportion of heterozygotes in the population decreases by a factor of 1 - *F*_*w*_, the variance in allele frequency increases with 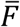, and the genetic diversity, 𝔼 = [2 *pq*], decreases as 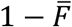.

Fig. S2B shows that in the absence of selection, the identity *F*_*b*_ increases slower than expected under the Wright–Fisher model with the actual population sizes (compare light shaded lines with black lines). These differences are a consequence of the circular mating scheme, which was designed to slow the loss of variation. The dotted line show the average *F*, estimated from the loss of heterozygosity in 50 replicate neutral simulations, each with 10^4^ loci on a chromosome of length *R*=1 Morgan. These are close to the prediction from the pedigree (light shaded lines), validating the simulations.

The thick colored line in fig. S2B show *F*, estimated in the same way from simulations that include truncation selection on a trait with within-family variance *V*_*s*_/*V*_*e*_ = 0.584 (a value we abbreviate as *θ* = 1), which matches the observed selection response and parent-offspring regression. The rate of drift, as measured by the gradient in *F* over time, is significantly faster in simulations with selection, by 6.7% in LS1 and 9.8% in LS2 (Student’s *t*-test *P* ≤ 0.008 in LS1 and *P* ≤ 0.0005 in LS2). However, this effect of selection would not be detectable from any one replicate, since the standard deviation of the rate of drift, relative to the mean rate, is ∼13% between replicates. On average, the observed loss of heterozygosity fits closely to that expected from the pedigree (large dot with error bars), though there is wide variation among chromosomes (filled dots), which is substantially higher than seen in simulations seeded with SNP at linkage equilibrium (compare filled and open dots).

We then performed 100 simulations, seeding each founding generation with actual genotypes and using actual pedigrees, selection pressure or heritability parameters (within-family heritability *h*^2^ of the fitness dimension: 0.51). A main conclusion from our modelling is that the overall allele frequencies were hardly perturbed by varying selection from random drift to even doubling the selection intensity. Upon closer examination, it became clear that under the standard “infinitesimal” model, selection could generate a weak but detectable excess of allele frequency sweeps compared to strict neutrality with no selection (fig. S2D, SNP classes 1/56 and 4/56). However, it would take many replicates (assuming no parallelism) for this excess to become statistically significant. Taken at face value, this result echoes many “evolve-and-resequence” (E&R) experiments based on diverse base populations that show only weak evidence of selective sweeps at specific loci (*23*, *75*).

#### Broader patterns and analyses of parallelism

On a broader scale, we also observed greater extent of parallelism globally than in the simulated results or with the empirical Ctrl line. For example, out of the 2405 and 2991 loci found above the *H*_*INF, no LD*_ cut-off in LS1 and LS2, 398 were found in both lines (13%; *χ*^2^ test, N∼150,000 windows; *χ*^2^=2901.4, d.f.=1, *P* ≤ 1 ×10^-10^); whereas we found only 10 or 7 overlaps in Ctrl–LS1 or Ctrl–LS2 comparisons, respectively. This difference is statistically significant (940 significant Ctrl loci at the *H*_*INF, no LD*_ threshold; N∼150,000 windows; Ctrl–LS1: *χ*^2^=0.7; Ctrl– LS2: *χ*^2^=6.0; both *P* = n.s.; see also fig. S8). In fact, there was not a single window out of a total of 8.4 million windows from the 100 replicates where both simulated LS1 and LS2 replicates simultaneously cleared the *H*_*INF, no LD*_ threshold. In contrast to our earlier analysis in single LS replicates, the parallel selected loci in both LS replicates loci may provide the strongest evidence yet to reject the infinitesimal model.

#### Enrichment for genes with functional impact on limb development

To determine what types of molecular changes may have mediated the selection response, we performed a gene set enrichment analysis. We asked if the outlier loci found in the Longshanks lines were enriched for genes affecting limb development (as indicated by their knockout phenotypes) and found increasingly significant enrichment as the allele frequency shift Δz^2^ cut-off became increasingly stringent (fig. S7A). The “limb/digital/tail” category of affected anatomical systems in the Mouse Genomic Informatics Gene Expression Database (*59*) showed the greatest excess of observed-to-expected ratio out of all 28 phenotype categories (the excluded “normal” category also showed no enrichment). In contrast, genes showing knock-out phenotypes in most other categories did not show similar enrichment as Δz^2^ became more stringent (fig. S7A). For genes expressed in limb tissue, there was a similar, but weaker increase, with the enrichment only appearing at higher Δz^2^ cut-off. We did not observe similar enrichment using data and thresholds derived from Ctrl (fig. S7A, lower panels). To investigate the impact on regulatory sequences, we obtained 21,211 limb enhancers predicted by ENCODE chromatin profile at a stage immediately preceding bone formation (Theiler Stage 23, at approximately embryonic day E14.5) (*57*). We found likewise an enrichment throughout the range of significance cut-offs (fig. S7A). Again, there was no similar enrichment in Ctrl.

#### Clustering with loci associated with human height

Since tibia lengths directly affect human height, we tested if an association exists between loci controlling human height (*64*) and a set of 810 loci at the *P* ≤ 0.05 significance level under *H*_*INF, no LD*_ described here. After remapping the human loci to their orthologous mouse positions (n = 655 out of 697 total height loci; data from GIANT Consortium), we detected significant clustering with the 810 peak loci (mean pairwise distance to remapped height loci: 1.41 Mbp vs. mean 1.69 Mbp from 1000 permutations of shuffled peak loci, range: 1.45–1.93 Mbp; n = 655 height loci and 810 peak loci; *P <* 0.001, permutations). We interpret this clustering to suggest that a shared and conserved genetic program exist between human height and tibia length and/or body mass.

#### Genome-wide analysis of the role of coding vs. cis-acting changes in response to selection

We examined the potential functional impact of coding or regulatory changes as a function of Δz^2^ in all three lines. For coding changes, we tracked the functional consequences of coding SNPs of moderate to high impact (missense mutations, gain or loss of stop codons, or frame-shifts). Whereas we found only mixed evidence of increased coding changes as Δz^2^ increased in the LS lines, there was a depletion of coding changes in Ctrl line as Δz^2^ increased, possibly due to purifying or background selection (fig. S7B; linear regression, LS1: *P* ≤ 0.015, slope > 0; LS2: *P* = 0.62, n.s., slope ≈ 0; Ctrl: *P* ≤ 5.72×10^-9^, slope < 0).

For regulatory changes, we used sequence conservation in limb enhancers overlapping a SNP as a proxy for functional impact. In contrast to the situation for coding changes, where the correlations differed between LS1 and LS2, the potential impact of regulatory changes increased significantly as a function of Δz^2^ in both LS lines (fig. S7B): within limb enhancers, SNP-flanking sequences became increasingly conserved at highly differentiated SNPs (phastCons conservation score, ranging from 0 to 1 for unconserved to completely conserved positions; linear regression, log-scale, *P <* 1.05×10^-9^ for both, slopes > 0). This relationship also exists for the Ctrl line, albeit principally from lower Δz^2^ and conservation values (*P <* 0.8×10^-3^, slope > 0; fig. S7B). Taken together, our enrichment analysis suggests that while both coding and regulatory changes were selected in the Longshanks experiment, the overall selection response may depend more consistently on *cis-*regulatory changes, especially for developmental regulators involved in limb, bone and/or cartilage development (Table 1; Table S3; c.f. Table S4 for coding changes). This is a key prediction of the “*cis-*regulatory hypothesis”, especially in its original scope on morphological traits (*76*).

#### Genes with amino acid changes of potentially major impact

We have further identified 12 candidate genes with likely functional impact on limb development due to specific amino acid changes showing large frequency shifts (albeit only one, *Fbn2*, cleared the stringent *P* ≤ 0.05 *H*_*INF, max LD*_ threshold; 6 in LS1, 9 in LS2, of which 3 were shared; Table S4). Consistent with strong selection for tibia development, all 12 genes show limb or tail phenotypes when knocked out, e.g., “short limbs” for the collagen gene *Col27a1* knockout. Most of these genes encode for structural cellular components, e.g., myosin, fibrillin and collagen (*Myo10; Fbn2*; and *Col27a1* respectively), with *Fuz* (fuzzy planar cell polarity protein) being the only classical developmental regulator gene. All but one of these genes have also been shown to have widespread pleiotropic effects with broad expression domains, and their knockouts were often lethal (eight out of 12) and/or exhibit defects in additional organ systems (11 out of 12). Based on this observation, we anticipate that the phenotypic impact of these selected coding missense SNPs (n.b. not knockout) would not be restricted to tibia or bone development.

#### Molecular dissection of Gli3, a candidate limb regulator, reveal gain-of-function cis-acting changes

We anticipated that genes related to major limb patterning, like *Gli3*, may contribute to the selection response (*77*, *78*). We thus performed an in-depth molecular dissection of *Gli3*, an important early limb developmental regulator on chromosome 13 (Chr13; Fig. S9A). This locus showed a substantial shift in minor allele frequency of up to 0.42 in LS1 (Δ*q*, 98^th^ quantile genome-wide, but below the *H*_*INF, max LD*_ threshold to qualify as a discrete major locus). We performed functional validation of *Gli3*, given its limb function (*79*) and considering that *Gli3* could be among the many minor loci in the polygenic background contributing to the selection response in LS1.

At the *Gli3* locus we could only find conservative amino acid changes (D1090E and I1326V) that are unlikely to impact protein function. Because the signal in LS1 was stronger in the 5’ flanking intergenic region, we examined the *Gli3 cis*-regulatory topologically associating domain (TADs, which mark chromosome segments with shared gene regulatory logic) (*22*) and identified putative enhancers using chromatin modification marks from the ENCODE project and our own ATAC-Seq data (Fig. S9B)(*28*, *57*). Four putative enhancers carried SNPs with large allele frequency changes. Among them, an upstream putative enhancer G2 (956 bp) carried 6 SNPs along with two 1-and 3-bp insertion/deletion (“indel”) with putative functional impact due to predicted gain or loss in transcription factor binding sites (Fig. S9C). We tested the G2 putative enhancer in a transgenic reporter assay by placing its sequence as a tandem duplicate upstream of a *lacZ* reporter gene (see Methods for details). We found that only the F17 LS1 allele was able to drive consistent *lacZ* expression in the developing limb buds (Fig. S9D). Importantly, this enhancer was active not only in the shaft of the limb bud but also in the anterior hand/foot plate, a major domain of *Gli3* expression and function (fig. S9A).

Furthermore, substitution of the enhancer sequence with the F0 allele (10 differences out of 956 or 960 bp) abolished *lacZ* expression (Fig. S9D). This showed that 10 or fewer changes within this novel enhancer sequence were sufficient to convert the inactive F0 allele into an active limb enhancer corresponding to the selected F17 allele (“gain-of-function”), suggesting that a standing genetic variant of the F17 allele may have been selectively favored because it drove stronger expression of *Gli3*, a gene essential for tibia development (*80*) [but see (*81*)].

#### Estimating the selection coefficient of the top-ranking locus, Nkx3-2, from changes in allele frequency

The significant locus on Chr5 containing *Nkx3-2* shows strong changes in SNP frequency in both LS1 and LS2. Here, we estimate the strength of selection on this locus, and the corresponding effect on the selected trait. We approximate by assuming two alternative alleles, and find the selection coefficient implied the observed parallel changes in allele frequency; we then set bounds on this estimate that take account of random drift. Finally, we use simulations that condition on the known pedigree to estimate the effect on the trait required to cause the observed strong frequency changes; these show that linked selection has little effect on the single-locus estimates.

We see strong and parallel changes in allele frequency at multiple steps. There are 14 non-overlapping 10kb windows that have a mean square change in arc-sin transformed allele frequency of 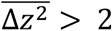 in both LS1 and LS2, spanning a 260 kbp region and including 807 SNP. SNP frequencies are tightly clustered, corresponding to two alternative haplotypes (Fig. 5A & fig. S11A). The initial (untransformed) allele frequencies average *q*_0_ = 0.18, 0.17 in LS1, LS2, respectively, and the final frequencies average *q*_17_ = 0.84, 0.98, respectively (also see fig. S11A, lower panel). These frequencies depend on the arbitrary threshold for which windows to include. However, this makes little difference, relative to the wide bounds on our estimates.

Under constant selection, 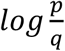 changes linearly with time, at a rate equal to the selection coefficient, *s*. Therefore, a naive estimate of selection is given by 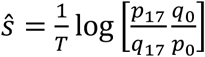 thus, ŝ =0.19, 0.32 for LS1, LS2, and averages 0.26. Here, males and females with longest legs are chosen to breed; the strength of selection on an additive allele depends on the fraction selected and the within-family trait variance. The former is kept constant, and there is little loss of variance due to inbreeding (*F*∼0.17), and so assuming constant selection is reasonable (Fig. 5B), unless there is strong dominance.

To set bounds on this estimate, we must account for random drift. The predicted loss of diversity over 17 generations, based on the pedigree, is *F*=0.173, 0.175 for LS1, LS2, which corresponds to an effective size *N*_*e*_ = 44.9, 44.4, respectively. Therefore, we calculate the matrix of transition probabilities for a Wright–Fisher population with 2*N* rounded to 90, 89 copies for LS1, LS2, over a range of selection, *s*. This yields the probability that the number of copies would change from the rounded values of 16/90 to 75/90 in LS1, and from 14/89 to 87/89 in LS2—that is, the likelihood of *s*, given the observed changes in allele frequency, and the known *N*_*e*_. There is no significant loss of likelihood by assuming the same selection in both lines; overall, ŝ = 0.24 (limits 0.13–0.36; fig. S11B).

#### Estimating the selection coefficient, accounting for linked loci

The estimates above using the simple approach do not account for selection on linked loci, and do not give the effect on the composite trait. We therefore simulated conditional on the pedigree and on the actual selection regime, as described above, but including an additive allele with effect *A* at the candidate locus on Chr5. The genetic variance associated with the unlinked infinitesimal background, and across Chr5, were reduced in proportion, to keep the overall heritability the same as before *V*_*a*_/(*V*_*a*_ + *V*_*e*_) = 0.539. The selection coefficient inferred from the simulated changes in allele frequency was approximately proportional to the effect on the trait, with best fit 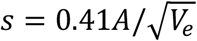 (fig. S11C, left). Assuming this relationship, we can compare the mean and standard deviation of allele frequency from simulations with linked selection, with that predicted by the single locus Wright–Fisher model (points vs. line in fig. S11C, middle & right). These agree well, showing that linked selection does not appreciably change the distribution of allele frequencies at a single locus. This is consistent with fig. S2D, which shows that linked selection only inflates the tail of the allele frequency distribution, an effect that would not be detectable at a single locus.

Combining our estimates of the selection coefficient with the relation 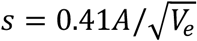, we estimate that the locus on Chr5 has effect 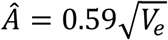, with 2-unit support limits 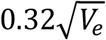 to 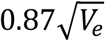. This single locus is responsible for ∼9.4% of the total selection response (limits 3.6– 15.5%).

This analysis does not allow for the inflation of effect that might arise from multiple testing. This is hard to estimate, because it depends on the distribution of effects across the genome, and also on the excess variation in estimates due to LD in the founder population. However, we note that if the effect of this locus is large enough that it would certainly be detected in this study, then there is no estimation bias from this source.

We also assume that there are two haplotypes, each with a definite effect. There might in fact be heterogeneity in the effects of each haplotype, for two reasons. First, this region might have had heterogeneous effects in the founder population, with multiple alleles at multiple causal loci. Second, as recombination breaks up the founder genomes, blocks of genome would become associated with different backgrounds. To the extent that genetic variation is spread evenly over an infinitesimal background, this latter effect is accounted for by our simulations, and has little consequence. However, we have not tested whether the data might be explained by more than two alleles, possibly at more than one discrete locus. Testing such complex models would be challenging, and we do not believe that such test would have much power. However, the estimates of selection made here should be regarded as effective values that may reflect a more complex reality.

**Table S1.** Sequencing Summary. For each line and generation, we individually barcoded all available individuals and pooled for sequencing. We aimed for a sequencing depth of around 100x coverage for 50–64 haplotypes per sample. Since the CD-1 mice were founded by an original import of 7 females and 2 males, we expect a maximum of 18 segregating haplotypes at any given locus. This sequencing design should give sufficient coverage to recover haplotypes genome-wide. Our successful genome-wide imputation results validated this strategy.

| Pool | n | Total reads | Mapped Sequence (Gbp) | Fold-coverage (x) | Median cov./Sample (x) |
| --- | --- | --- | --- | --- | --- |
| Ctrl, F0 | 25 | 1,856,046,931 | 251.3 | 92.0 | 2.42 |
| LS1, F0 | 26 | 1,858,953,260 | 256.1 | 93.8 | 2.82 |
| LS2, F0 | 25 | 2,011,646,609 | 283.2 | 103.7 | 3.57 |
| Ctrl, F17 | 32 | 1,882,838,451 | 260.2 | 95.3 | 2.97 |
| LS1, F17 | 32 | 2,071,122,164 | 292.1 | 107.0 | 2.95 |
| LS2, F17 | 31 | 1,897,174,855 | 267.0 | 97.8 | 2.93 |
| **Total** | **169** | **11,577,782,270** | **1609.9** | **589.5** | **2.91** |

**Table S2.** Pairwise F_ST_ and segregating sites (S) between populations. As expected, there is a general trend of decrease in diversity after 17 generations of breeding. Globally, there was a 13% decrease in diversity, but F17 populations still retained on average ∼5.8M segregating SNPs (diagonal). There was very little population differentiation, as indicated by low F_ST_ among the three founder populations, however F_ST_ increases by 100-fold among lines by generation F17 (above diagonal, orange boxes). Within-line F_ST_ is intermediate in this respect, increasing by about half that amount.

| S \ F<sub>ST</sub> | Ctrl F0 | LS1 F0 | LS2 F0 | Ctrl F17 | LS1 F17 | LS2 F17 |
| --- | --- | --- | --- | --- | --- | --- |
| Ctrl F0 | 7,218,921 | 0.00095 | 0.00176 | 0.06067 | 0.09601 | 0.10694 |
| LS1 F0 | 7,213,186 | 6,612,653 | 0.00100 | 0.07731 | 0.08045 | 0.10830 |
| LS2 F0 | 7,306,904 | 7,282,910 | 7,239,774 | 0.07535 | 0.09505 | 0.09634 |
| Ctrl F17 | 7,101,376 | 7,118,072 | 7,227,135 | 5,847,708 | 0.17469 | 0.17942 |
| LS1 F17 | 7,121,979 | 7,062,183 | 7,214,508 | 6,795,924 | 5,813,162 | 0.17631 |
| LS2 F17 | 7,218,921 | 7,155,580 | 7,239,774 | 6,832,220 | 6,778,532 | 5,749,742 |

**Table S3.**
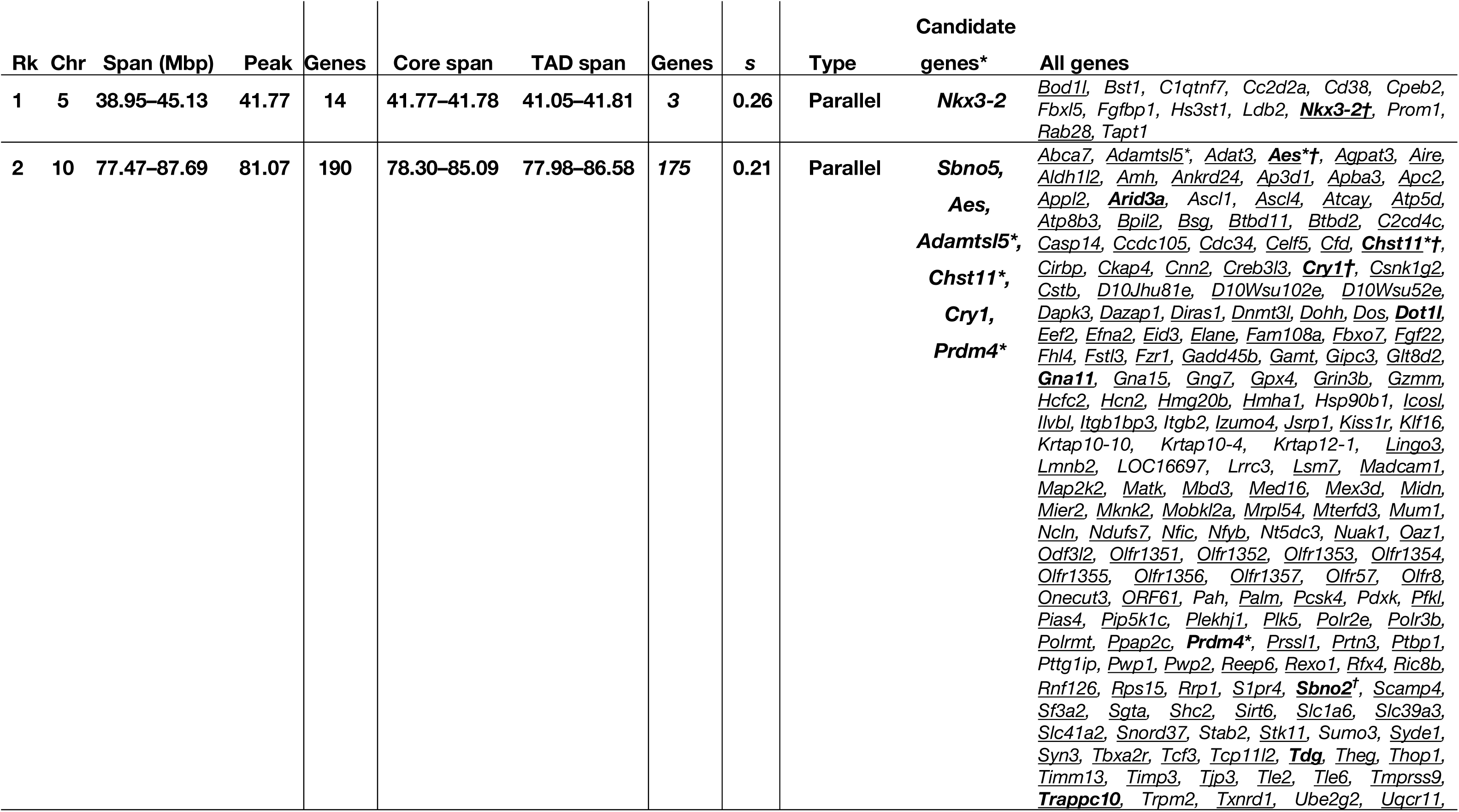
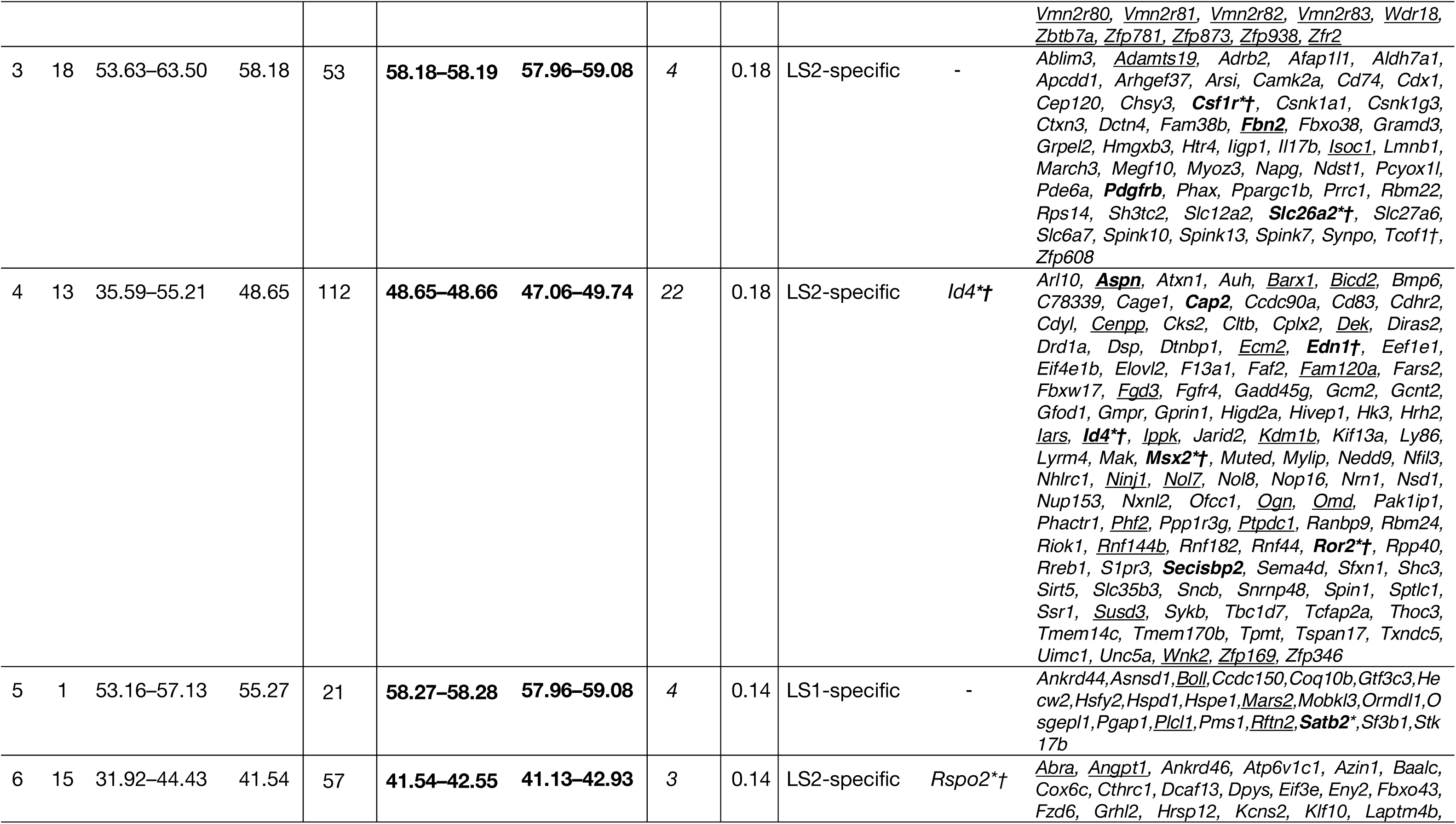
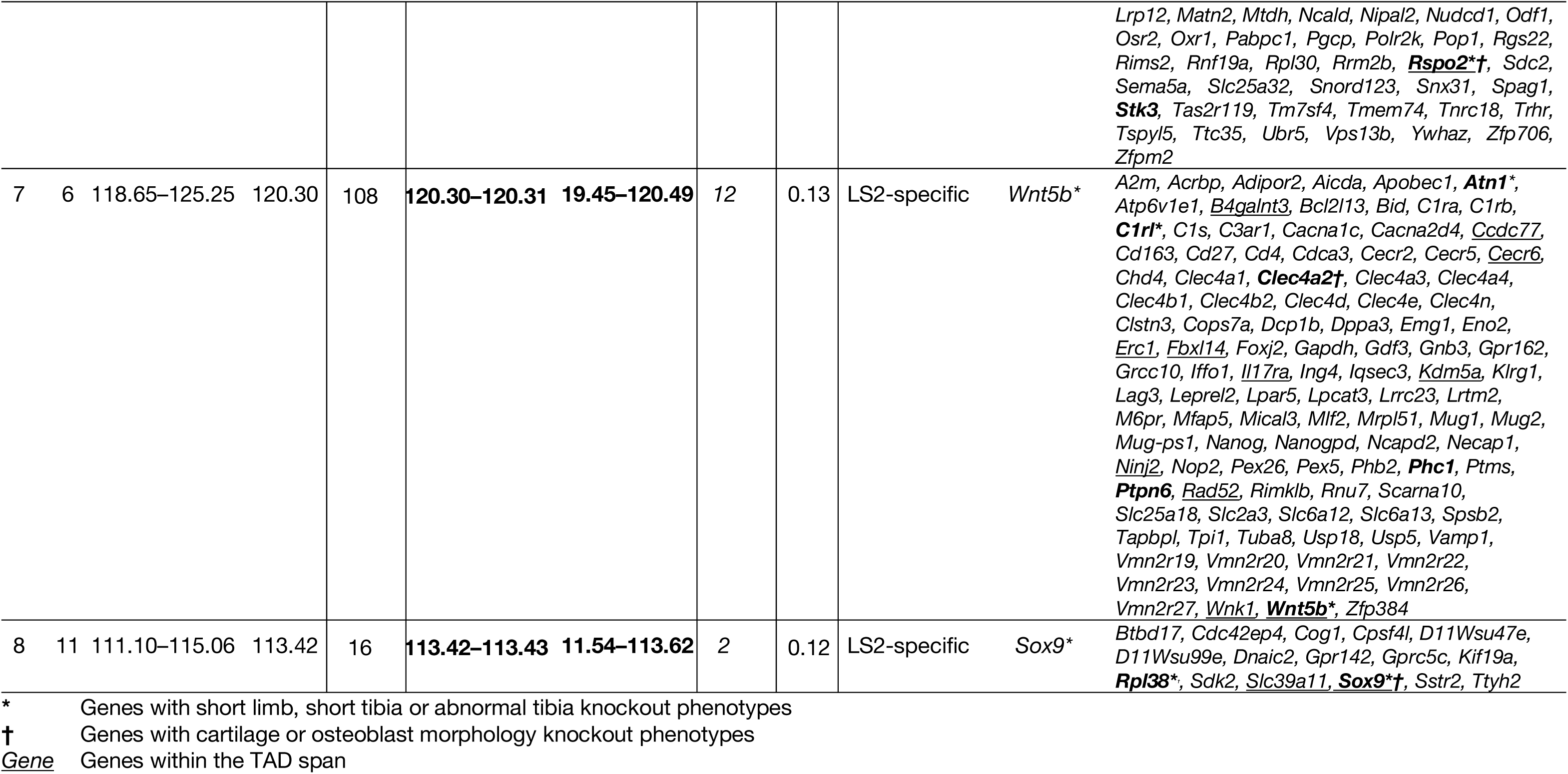
Full details on the eight discrete loci. Listed here are the eight loci shown in Table 1, with additional details on the core span and the TAD span used to identify candidate genes, and a full list of genes within the full span.

**Table S4.**
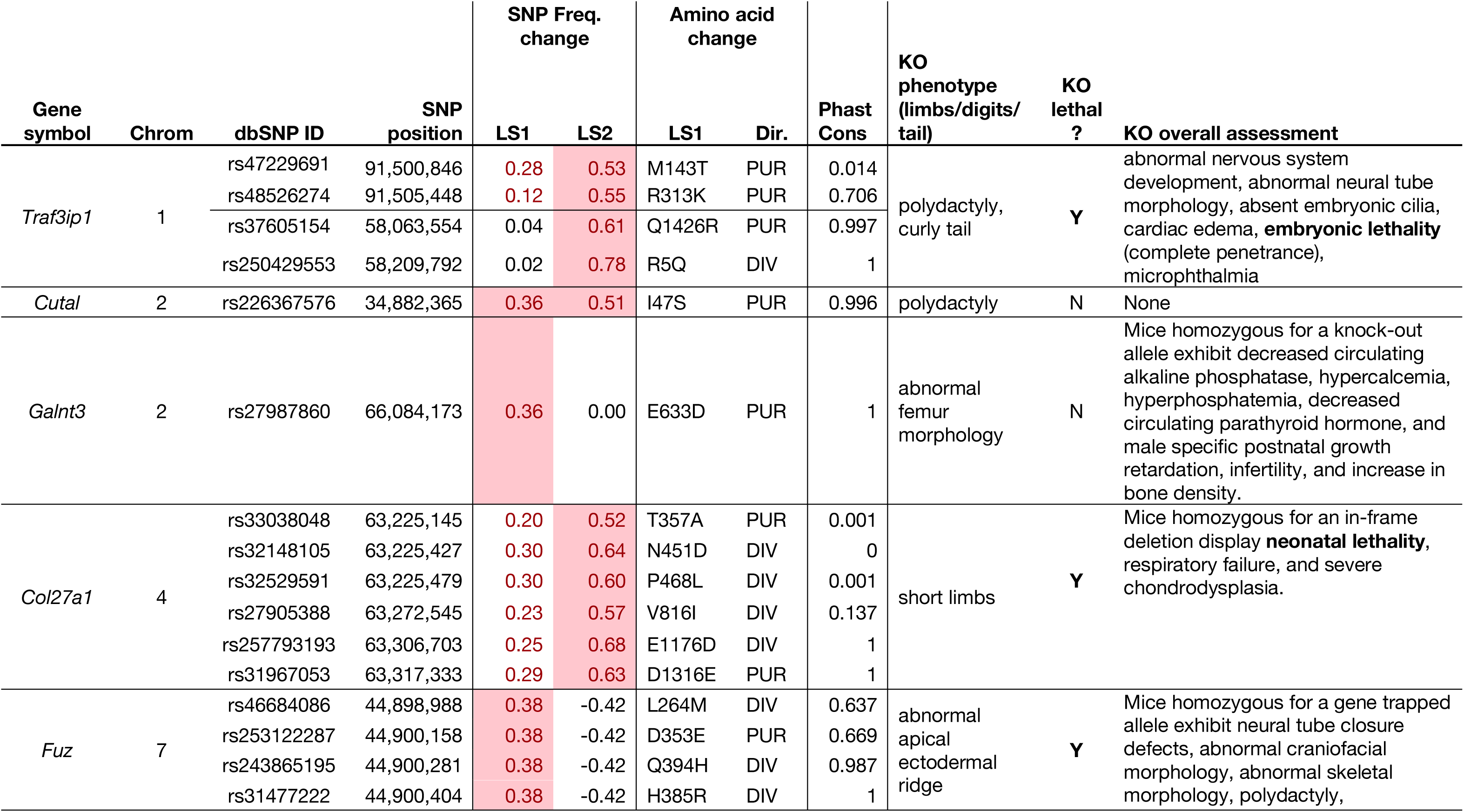
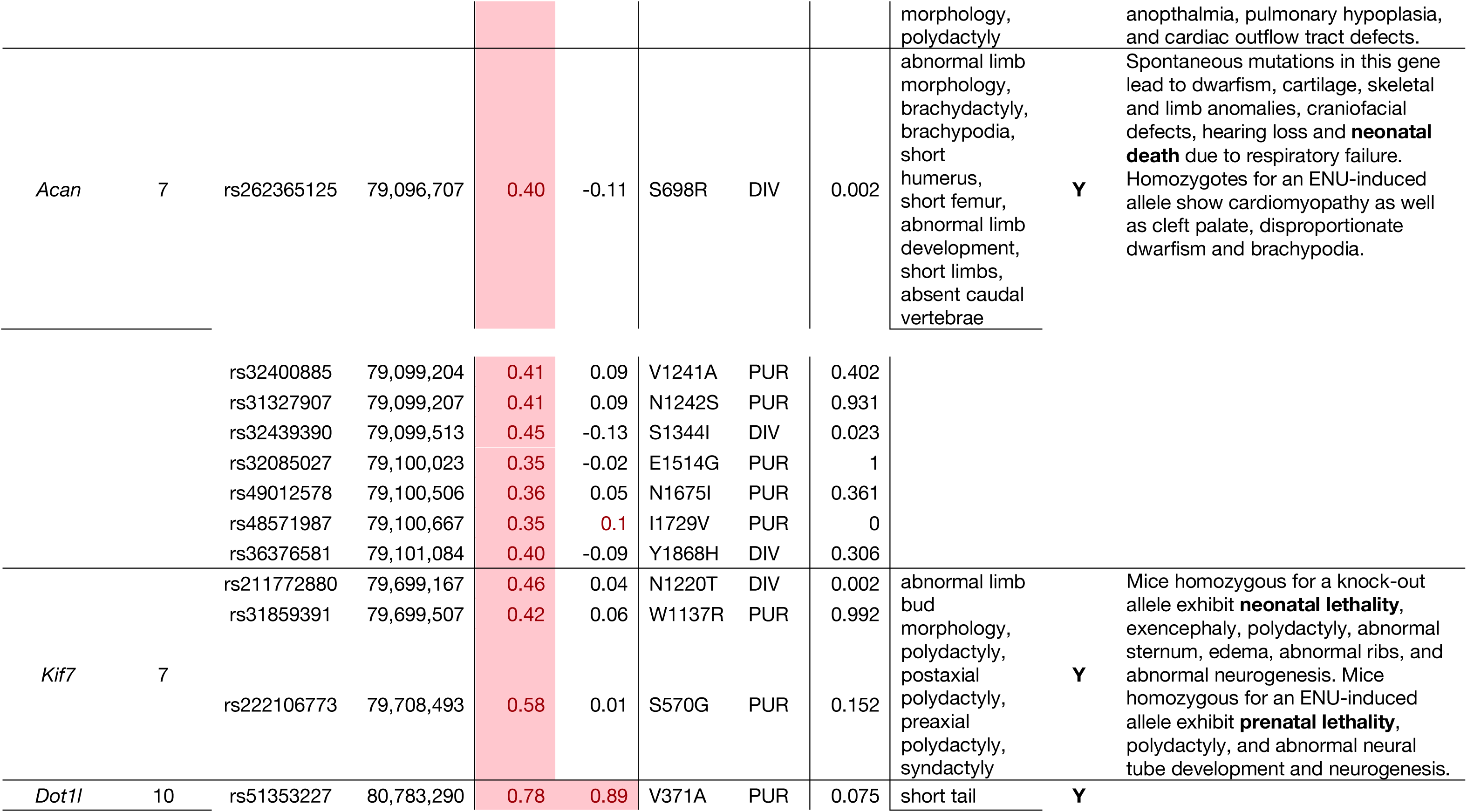
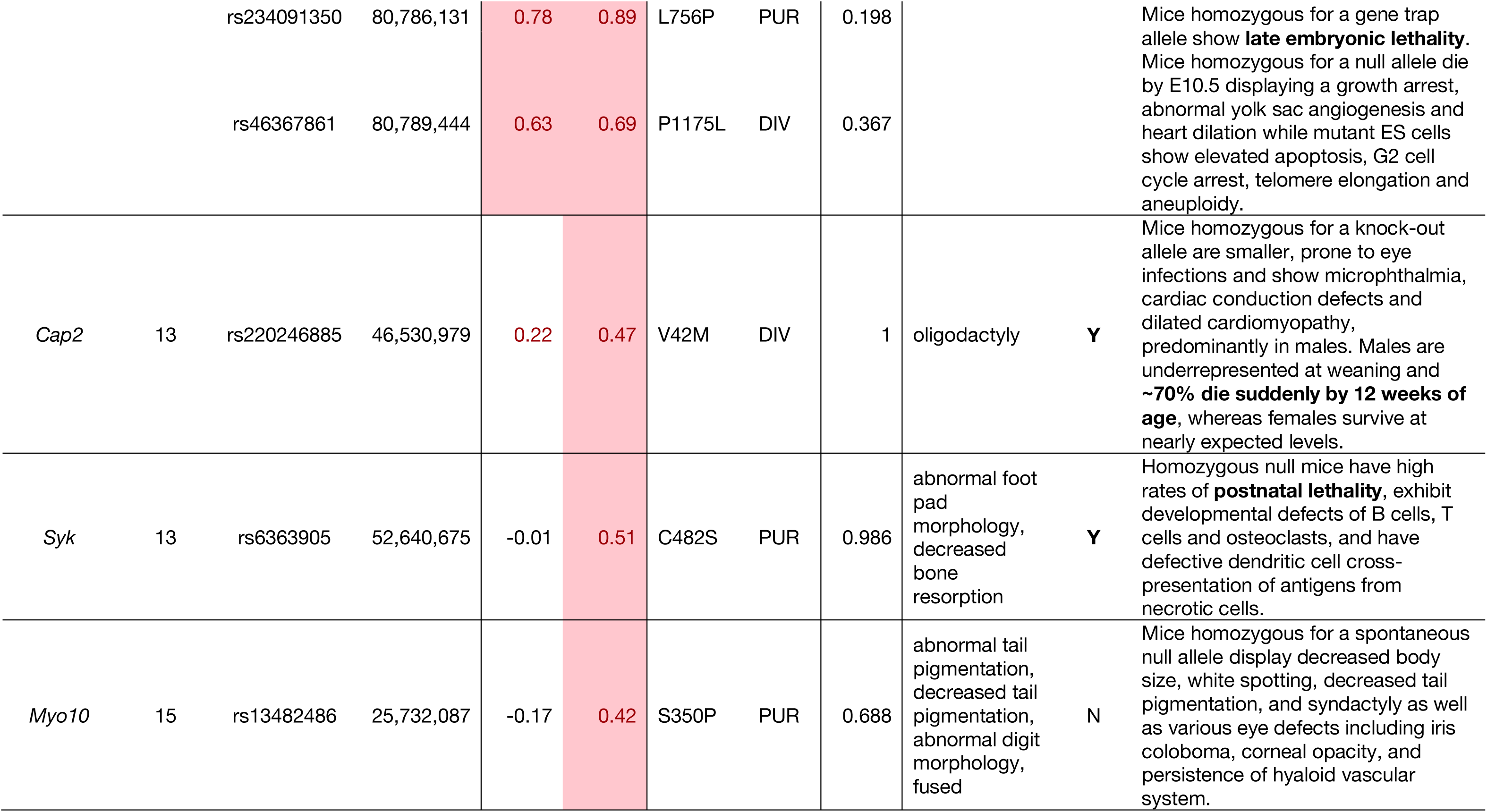
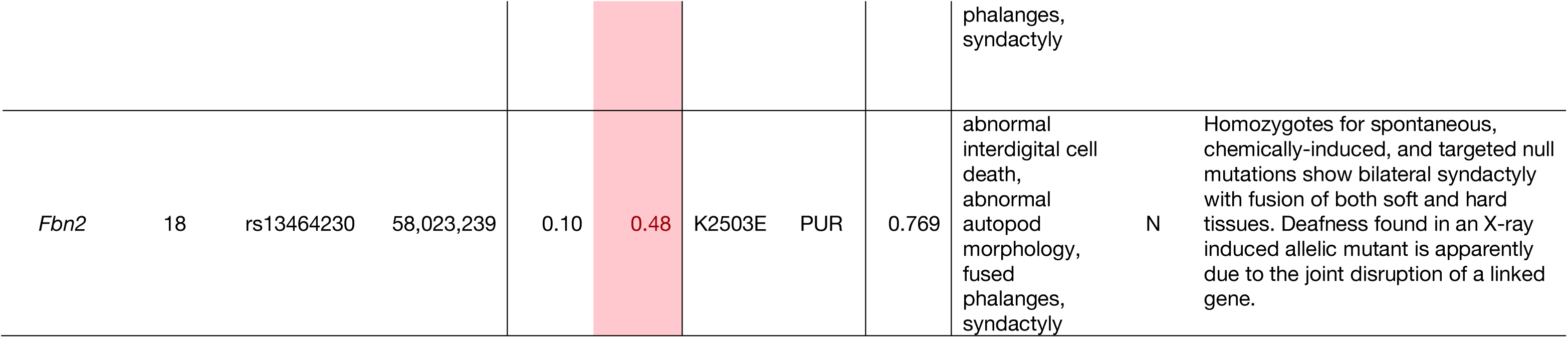
Detected protein-coding changes with large allele frequency shift in amino acids. Listed are genes carrying large frequency changing SNPs affecting amino acid residues. Highlighted cells indicate the line with greater frequency changes ≥ 0.34 (red text with shading). Other suggestive changes are also shown with red numbers in unshaded cells. The changed amino acids are marked using standard notations, with the directionality indicated as “purifying” or “diversifying” with respect to a 60-way protein sequence alignment with other placental mammals. The conservation score based on phastCons was calculated at the SNP position itself, ranging from 0 (no conservation) to 1 (complete conservation) among the 60 placental mammals. For each gene, reported knockout phenotypes of the “limbs/digits/tail” category is reported, along with whether lethality was reported in any of the alleles, excluding compound genotypes. A summary of the mutant phenotype as reported by the Mouse Genome Informatics database is also included to highlight any systemic defects beyond the “limbs/digits/tail” category (lethal phenotypes reported in **bold**).

**Table S5.**
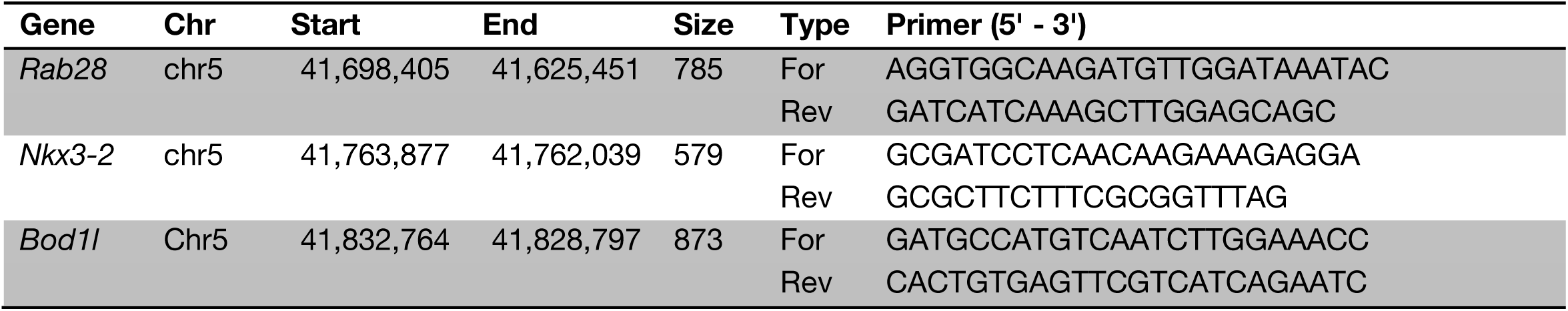
Oligonucleotides for *in situ* hybridization probes. VP: Viewpoint.

**Table S6.**
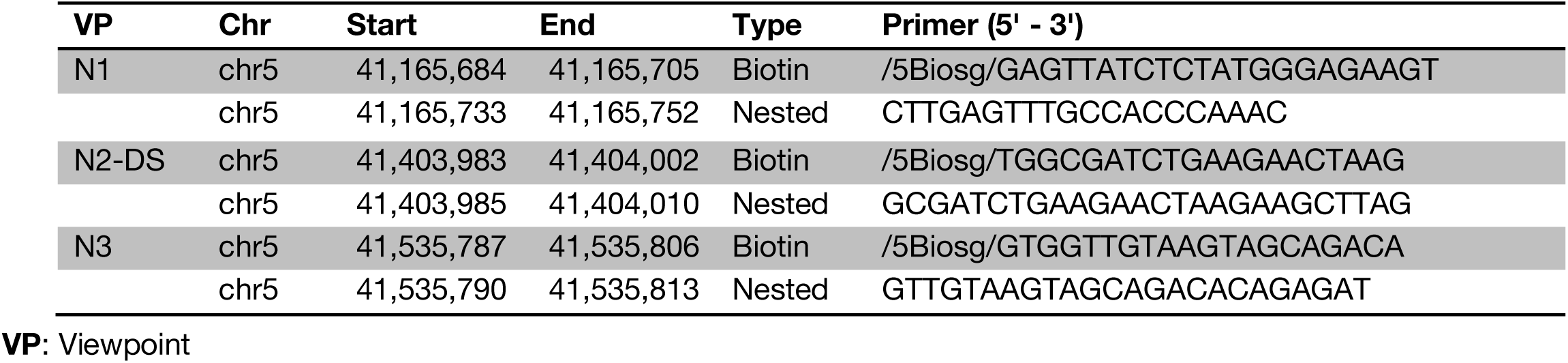
Oligonucleotide primers for multiplexed 4C-seq of enhancer viewpoints at the *Nkx3-2* locus. The 4C-seq adapter and adapter-specific primer sequences are given in (*69*). N2-DS denotes its location as 18 kbp downstream of the actual N2 enhancer. All viewpoints are pointed towards *Nkx3-2* gene body (“+” strand). **Enh**: Enhancer

**Table S7.**
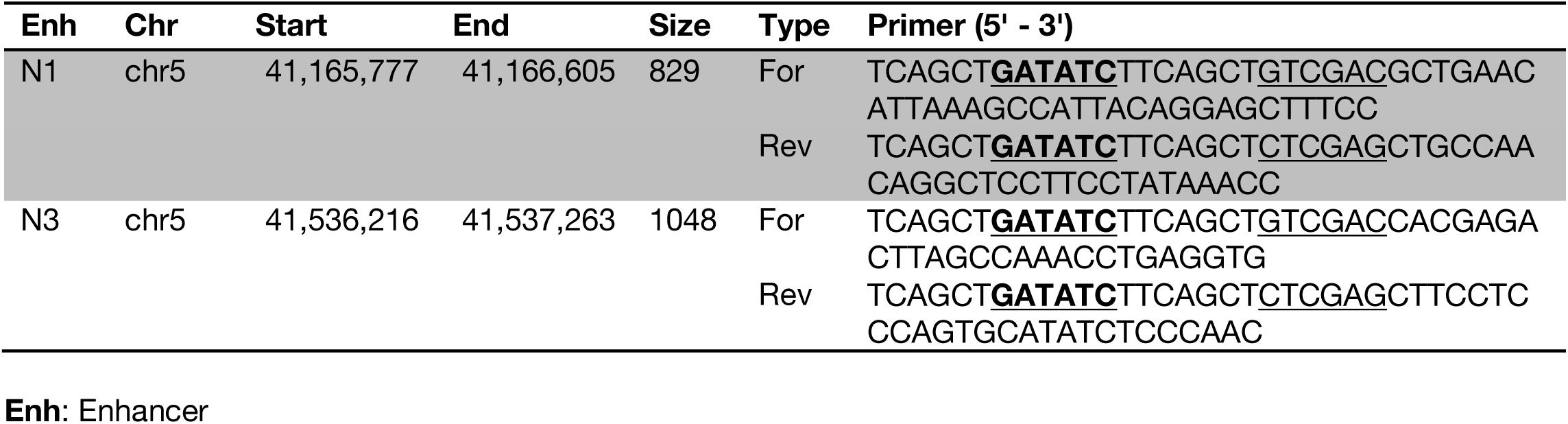
Oligonucleotide primers for amplifying the enhancers at the *Nkx3-2* locus. Each of the amplicons are tagged with *SalI* (forward) or *XhoI* (reverse) sites (underlined) for concatenation and flanked by *EcoRV* sites (underlined and bold) for insertion into the pBeta-lacZ-attBx2 reporter vector upstream of the β-globin minimal promoter.

**Table S8.**
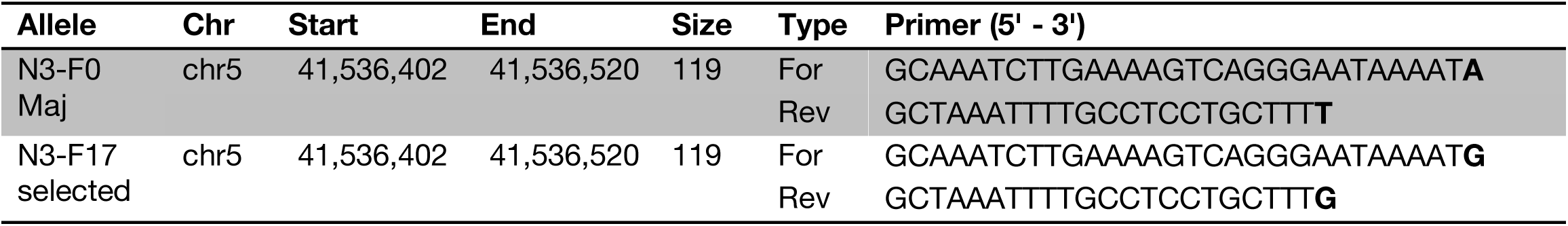
Oligonucleotide primers for allele-specific genotyping of the N3 enhancer. The primers were designed to target two SNPs (bold) in the N3 enhancer, rs33219710 and rs33600994.

**fig. S1.**
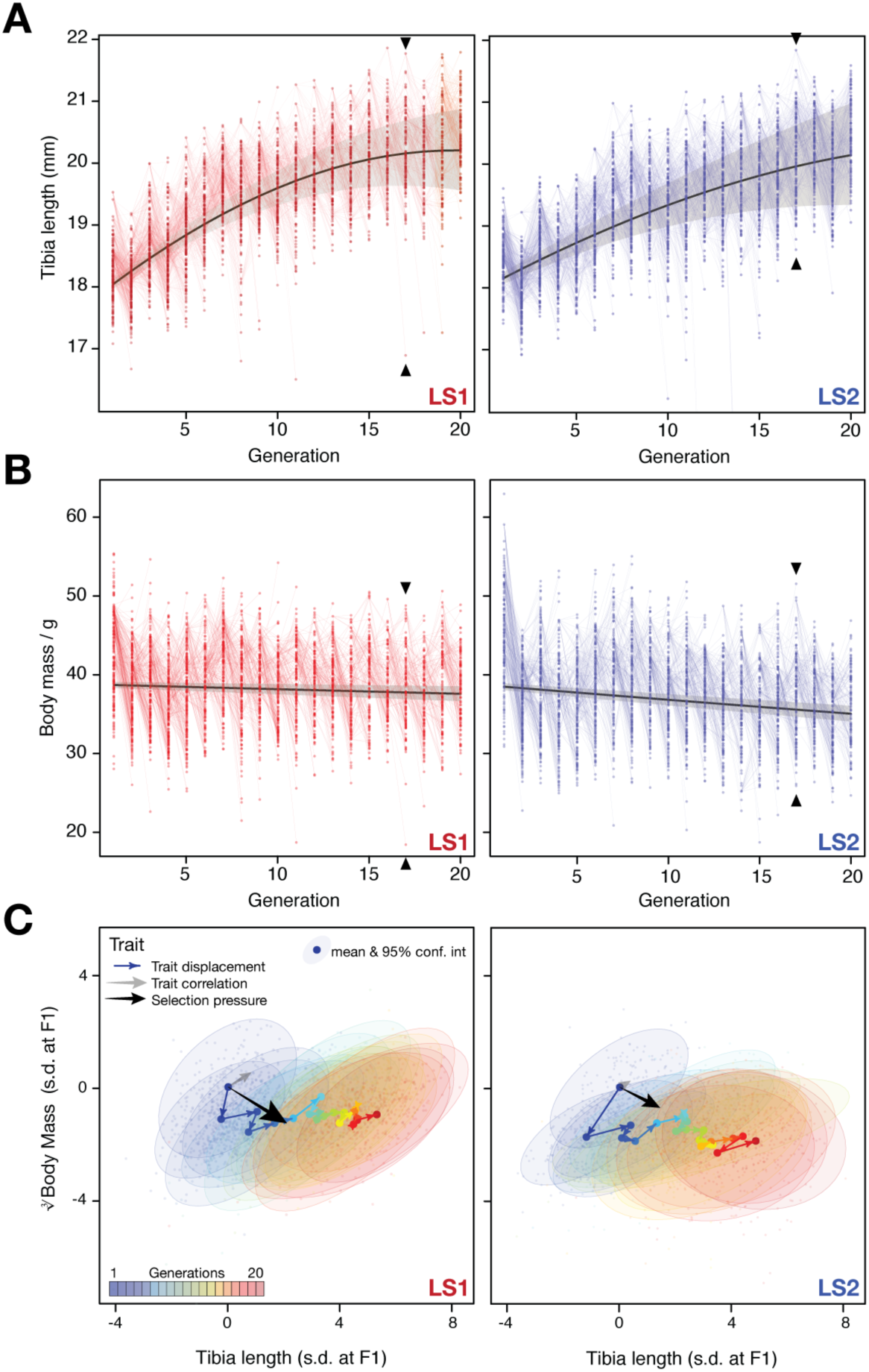
Artificial selection allowed detailed reconstruction of selection parameters. Rapid response to selection produced mice with progressively longer tibiae (**A**) and slightly lower body weight (**B**) within 20 generations. Having complete records throughout the selection experiment makes it possible to reconstruct the selection response for both phenotypes and genotypes in detail. Individuals varied in tibia length in both Longshanks lines (LS1, left; LS2, right). Lines connect parents to their offspring. The actual selection depended on the within-family and within-sex rank order of the tibia length-to-body mass (cube root) ratio (see (*14*) for details). The overall selection response was immediate and rapid for tibia length (**A**), suggesting a selection response that depended on standing variation among the founders (black lines show the best fitting quadratic function, with shading indicating 95% confidence interval; adjusted R^2^ = 0.61 for LS1; 0.43 for LS2). Strong selection response led to rapid increase in tibia length. In contrast, there was only minor decrease in body weight over the course of the experiment. (**C**) Trajectory in selection response shows decoupling of correlation between tibia length and body mass. Despite overall correlation between tibia length and body mass (grey arrow and major axes in confidence envelopes), cumulative trait displacement over the 20 generations (expressed in s. d. units at F1; arrows, dots and 95% confidence envelopes, color-coded according to generation) showed persistent increase in tibia length with only minor change in body mass along the general direction of selection pressure (black arrows from F1; vector length and directions based on logistic regression). This shows that the Longshanks selection experiment was successful in specifically selecting for increased tibia length while keeping relatively unchanged body mass.

**fig. S2.**
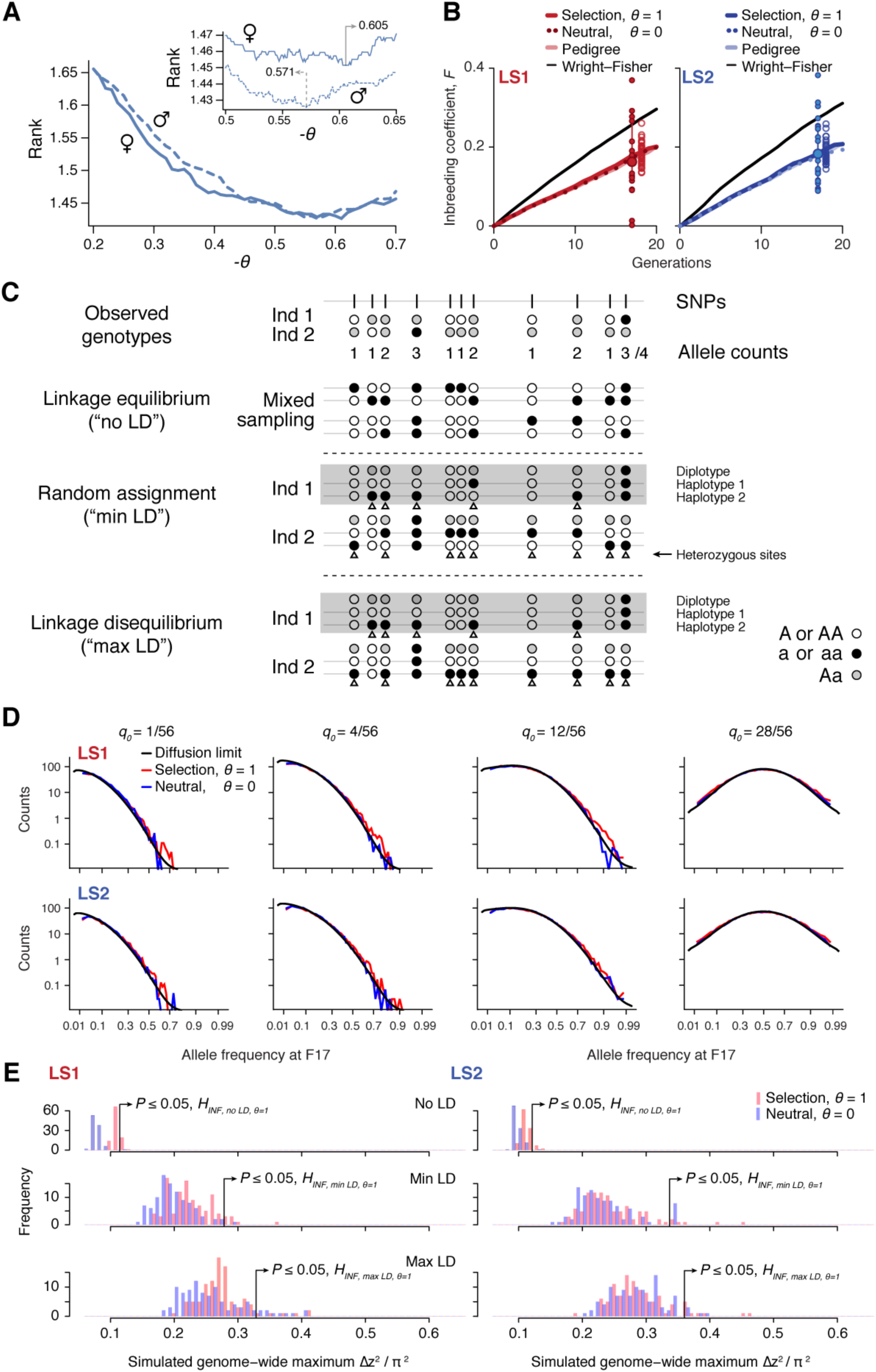
Simulating selection on pedigrees. (**A**) Finding the correct (value for the composite trait *in*(***TB***^θ^). In each simulated family, offsprings are split by sex and ranked by their composite trait. Due to occasional use of back-up crosses, the average rank of actual breeders is greater than 1. We vary (to find the value where actual breeders in the LS lines have the best (lowest) rank. We find (= -0.571 to show the best match for males and 0.605 for females. For subsequent analyses we set (to be -0.57. (**B**) Increase in inbreeding over the course of the Longshanks experiment. The lines show the change in identity between two alleles between diploid individuals, *F*_*b*_, over 20 generations, as calculated from the pedigree (light shade); an average of 50 neutral simulations without selection (dotted line); or the average of 50 simulation replicates with selection intensity at *V*_*s*_/*V*_*e*_ = 0.584 (*θ* = 1; thick, dark line). While the *F*_*b*_ trajectories based on pedigree or neutral simulations are indistinguishable, inbreeding increases slightly faster under selection (thick line). The black line shows the increase in identity expected under a Wright–Fisher model with the actual population sizes; under this model, *F*_*w*_ and *F*_*b*_ are close to each other, and to 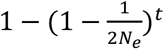, with *N*_*e*_ equal to the harmonic mean, 24.8. The large dot (with error bar showing the interquartile range among chromosomes) at right show the actual *F*_*b*_, estimated from the decline in average 2*p* (1 - *p*) over 17 generations. Small filled dots show the estimates from each of the 20 chromosome. Open dots show 40 replicate simulations, made with the same pedigree and the same selection response (= 1 and sub-sampling from the simulated chromosome according to the actual map length of each of the mouse chromosomes (*46*). The simulation agrees well with the observed genome-wide average. Most of the observed data from chromosomes fall within a range comparable to simulated replicates (compare large dot with open dots), with LD being the likely source of this excess variance. (**C**) Three different schemes to seed founder haplotypes. We simulate founder haplotypes that are consistent with observed genotypes (shown here as black, white and grey dots as the two homozygous and the heterozygous states) by directly sampling from founder individuals in each LS line. Under the linkage equilibrium scheme, we sample from the list of allele counts at all SNPs. This produces founder haplotypes that carry no linkage disequilibrium (“no LD”). Under the random assignment scheme, we sample according to each individual (shown as “diplotypes” within the box for easy comparison). At heterozygote sites in each individual (arrowheads), we randomly assign the alleles to the two haplotypes. This produces founder haplotypes that show minimal LD that is consistent with the observed genotypes (“min LD”). Under the “max LD” assignment scheme, we also sample according to each individual, except that we consistently assign its haplotypes 1 and 2 with reference (white) and alternate (black) alleles, respectively. This maximizes LD in the founder haplotypes (“max LD”). (**D**) Simulated vs. expected allele frequency shifts. The distribution of allele frequencies at generation 17 is compared with the distribution expected with no selection (blue) or with selection (red), given a frequency of 1, 4, 12 or 28 minor alleles out of 56 founding alleles. The black line shows the diffusion limit, calculated for scaled time 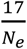, with *N*_*e*_ estimated to be 51.7 and 48 in LS1 and LS2 respectively, from the rate of increase in *F*, calculated from the pedigree in panel **A** above. (**E**) Significance threshold values under varying LD from 100 simulated replicates (blue: no selection; red: observed selection response in the actual experiment, (= 1; see panel **C** on LD assignment methods). In order to account for non-independence of adjacent windows due to linkage, a distribution of genome-wide *maximum* Δz^2^ was used to determine the significance threshold at each LD level. As seen in previous panels, increasing selection pressure does produce greater shifts in Δz^2^ despite using the same pedigree due to a relatively greater proportion of additive genetic variance *V*_*s*_. However, a far greater impact on Δz^2^ is due to changes in LD. This is because weak associations between large numbers of SNP can greatly inflate the variance of Δz^2^. Of the three LD levels, “max LD” likely produced overly conservative thresholds, whereas “min LD” may lead to higher false positives. We have opted conservatively to use maximal LD in our analysis.

**fig. S3.**
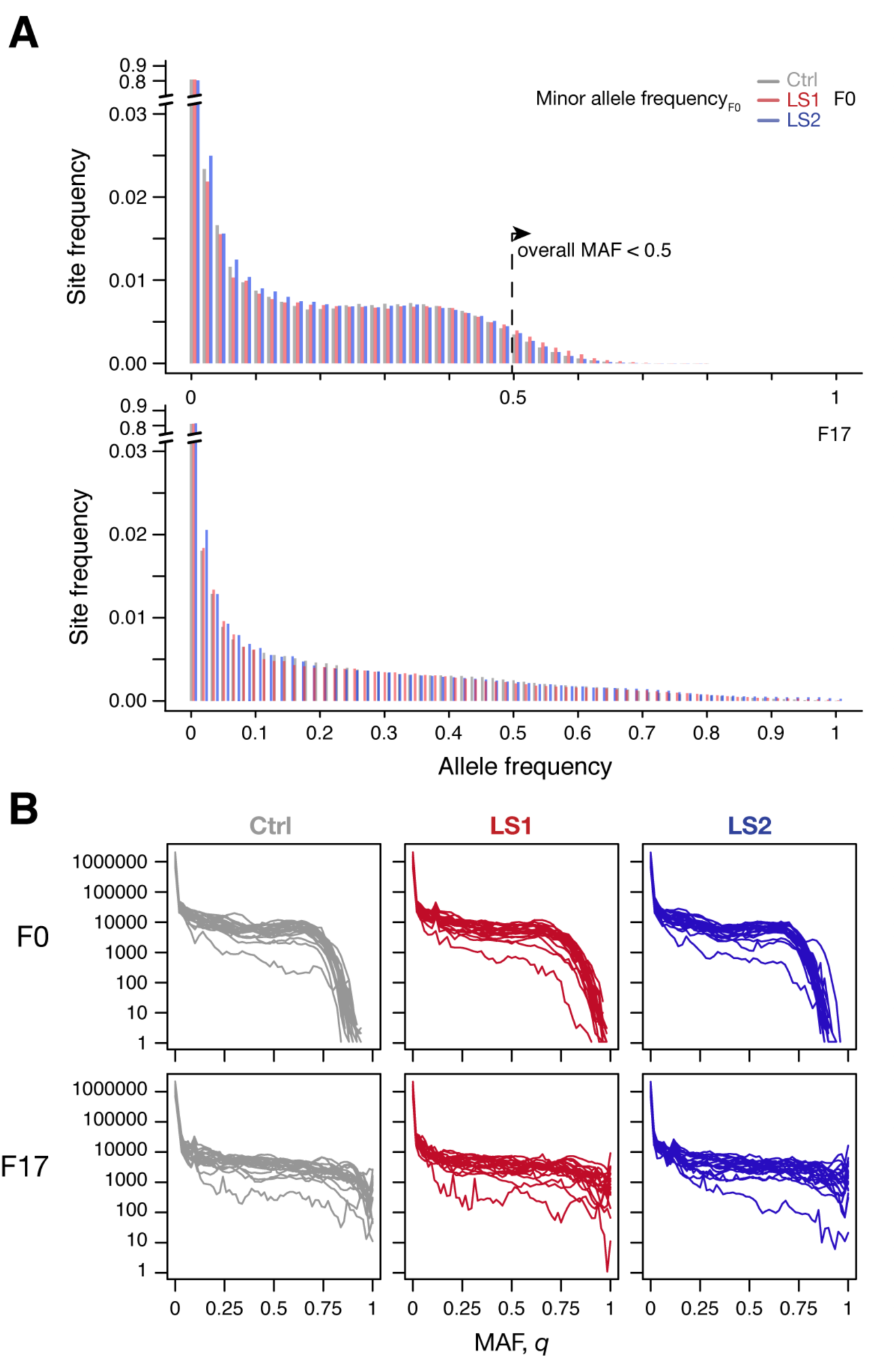
Broad similarity in molecular diversity in the founder populations for the Longshanks lines and the Control line. (**A**) Shown are the site frequency spectra from LS1, LS2 and Control lines at F0 (top, folded, based on a global minor allele frequency or MAF ≤ 0.5) and F17 (bottom, unfolded, but tracking the same minor allele as in F0). Overall the spectra were very similar to each other. The Control population was mostly intermediate in the decay in the rarer alleles. A small number of sites show MAF > 0.5 in each line, even though the overall MAF is ≤ 0.5. After 17 generations, the same alleles were generally more spread out, leading to more broadly distributed spectra. There was again little overall change between the Longshanks and Control lines. Due to differing number of available haplotypes, the F17 bars may appear to change in their grouping. (**B**) Variations between chromosomes shown in each line and generation. The unfolded site frequency spectrum is shown based on the MAF assigned as in **A**. There is substantial variation between chromosomes, which shows increased distortions in F17.

**fig. S4.**
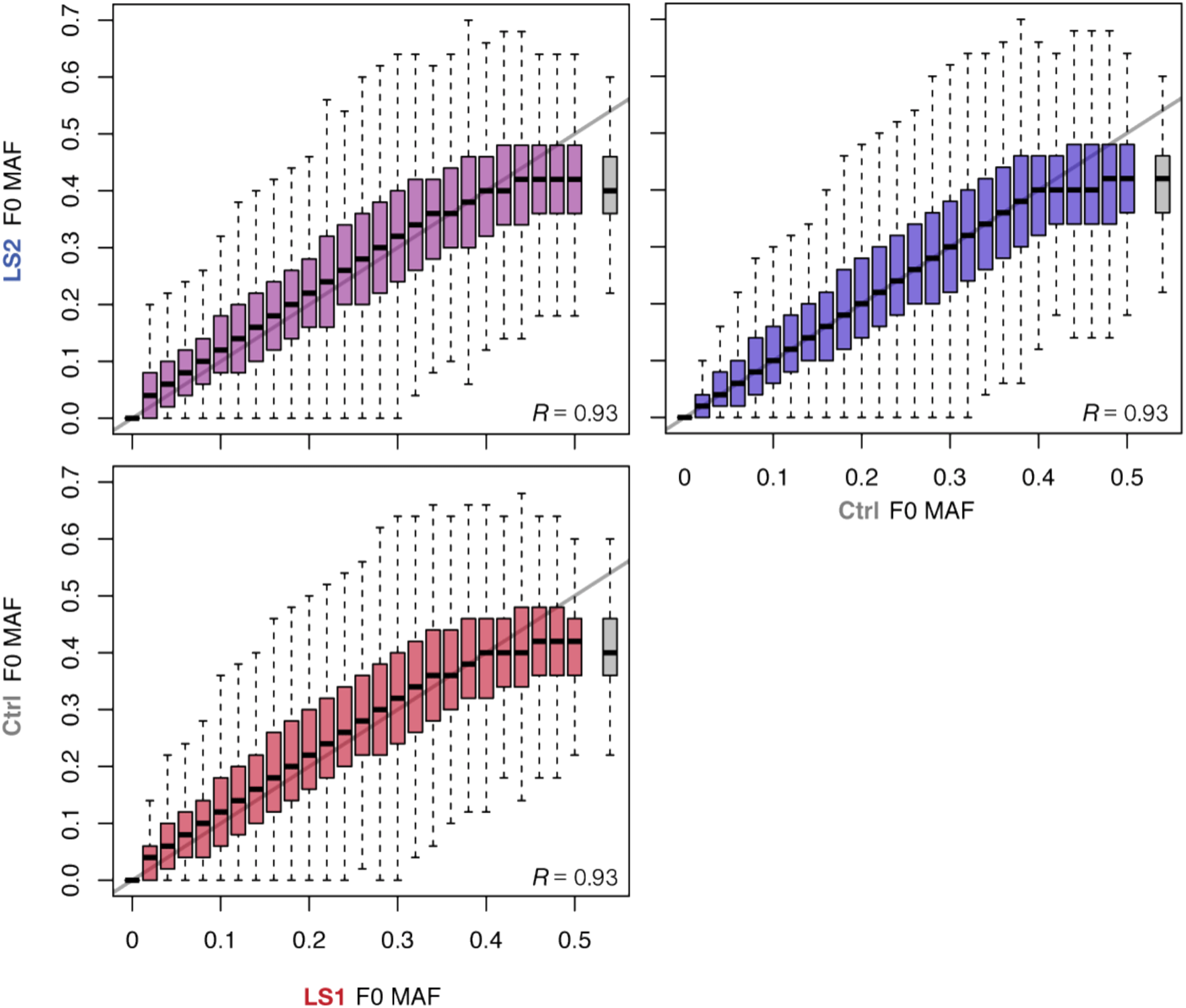
Allele frequencies between the founder populations were very similar. Joint minor allele frequencies shown as box plots in 2% bands between the Control and LS1 (red), LS2 (blue); or the two LS lines (purple). Outliers were omitted for clarity. The overall trends follow closely the parity line (grey line along the diagonal), except at frequencies very close to 0.5. Similar to the site frequency spectra shown in fig. S3A, a small number of sites have a MAF above 0.5 (grey box), because of the use of an overall MAF ≤ 0.5 to determine minor allele status to enable comparisons across lines. Correlations between all pairwise combinations were around 0.93.

**figure S5.**
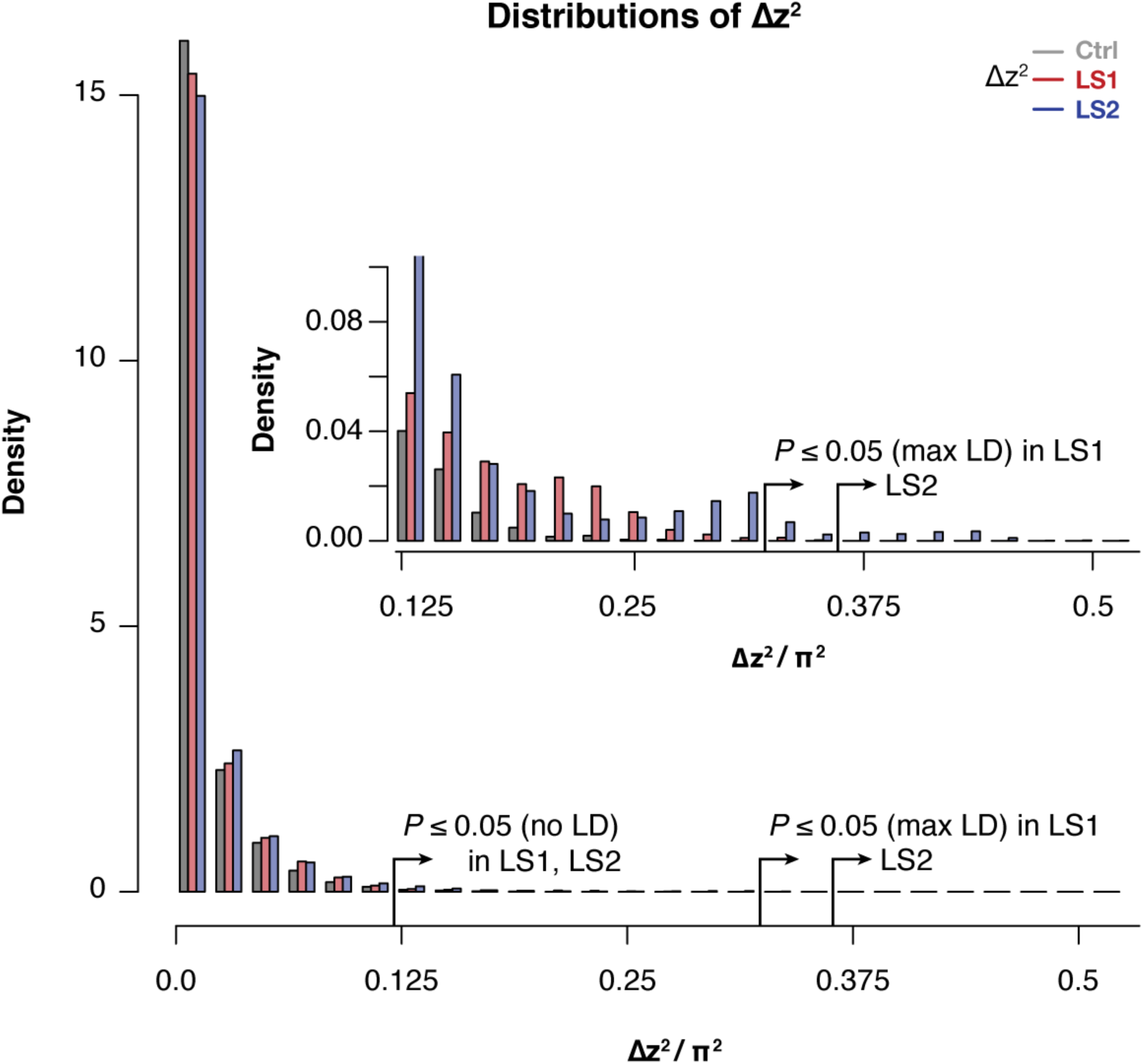
Selected lines showed more extreme values of Δz^2^ than the Control line. Histogram of within-line Δz^2^ values in 10 kbp windows across the genome in the LS1, LS2 and Control. Overall similarity is high across all 3 lines, but there was an excess of large Δz^2^ value starting from as low as < 0.1 π^2^. This pattern becomes clearly distinct above the threshold value of 0.125, which corresponds to the lenient significance threshold *P* ≤ 0.05 under *H*_*INF,* *no* *LD*_ (inset). There were clearly an excess of windows in LS2 above the more stringent *P* ≤ 0.05 threshold under *H*_*INF,* *max* *LD*_. Such excess supports discrete loci contributing to selection response in LS2 that give rise to greater distortion of Δz^2^ spectra.

**fig. S6.**
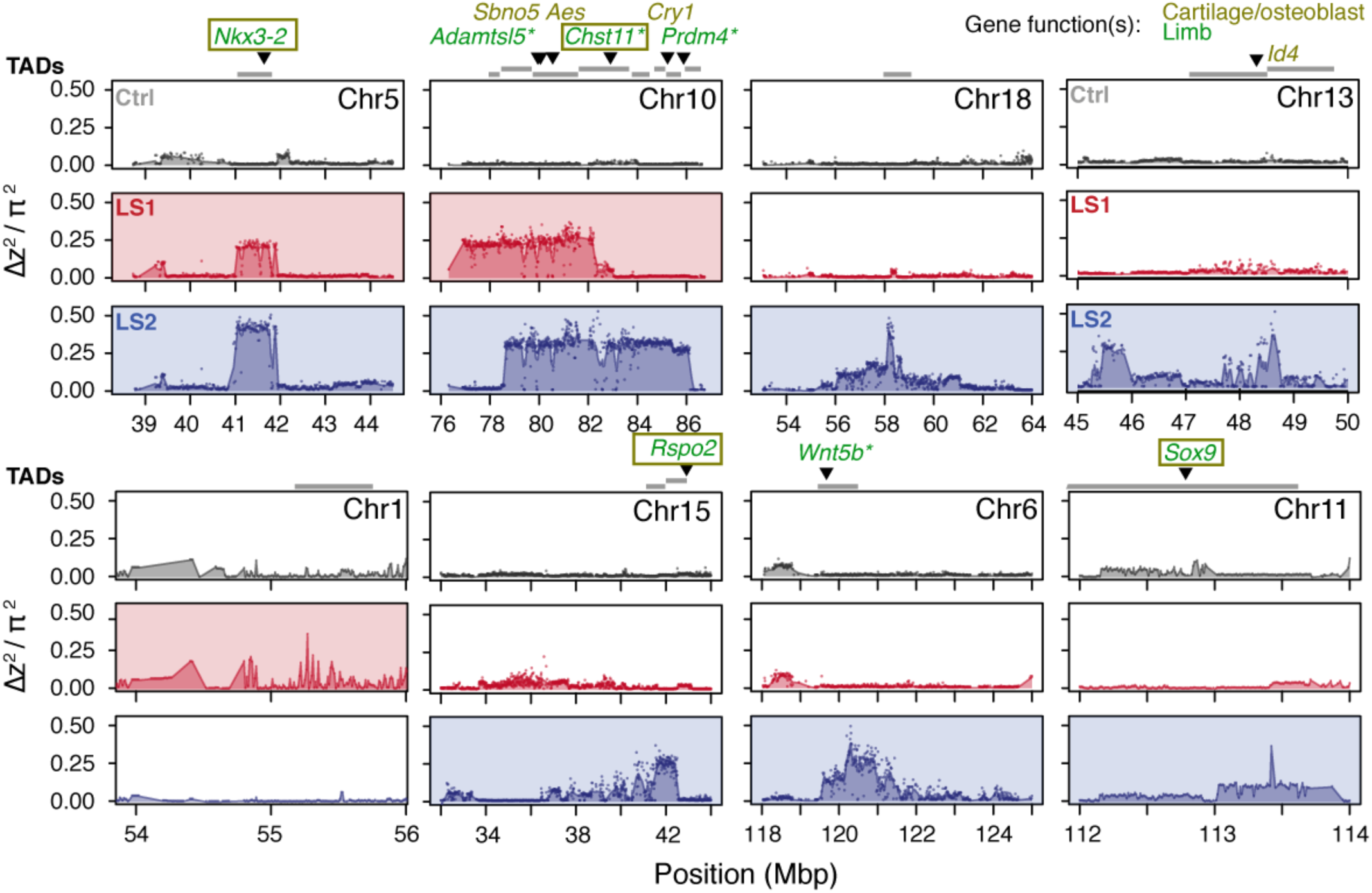
Detailed Δz^2^ profiles at the LS significant loci. For each significant locus, Δz^2^ profiles are shown for Ctrl (grey), LS1 (red) and LS2 (blue). Plots are shaded if the locus is significant in a given line. TADs within 250 kbp of the significant signals are shown as grey bars above each locus, with gene whose knockout phenotypes of the following categories highlighted: “abnormal tibia morphology”, “short limb”, “short tibia”, “abnormal cartilage morphology”, “abnormal osteoblast morphology”. The gene symbols are colored according to the gene function(s) in limb development (green), in bone development (yellow) or both (boxed). Gene symbols marked by asterisk (*) have reported “short tibia” or “short limb” knockout phenotypes. All of the above categories show significant enrichment at the 8 loci (number of genes per category: 4–7, nominal *P* ≤ 0.03, see Supplementary Methods, section *Candidate genes* for details on the permutation), except “abnormal cartilage morphology”, with 4 genes and a nominal *P*-value of 0.083. No overlap was found with any gene in these categories from the three significant loci from the Ctrl line.

**fig. S7.**
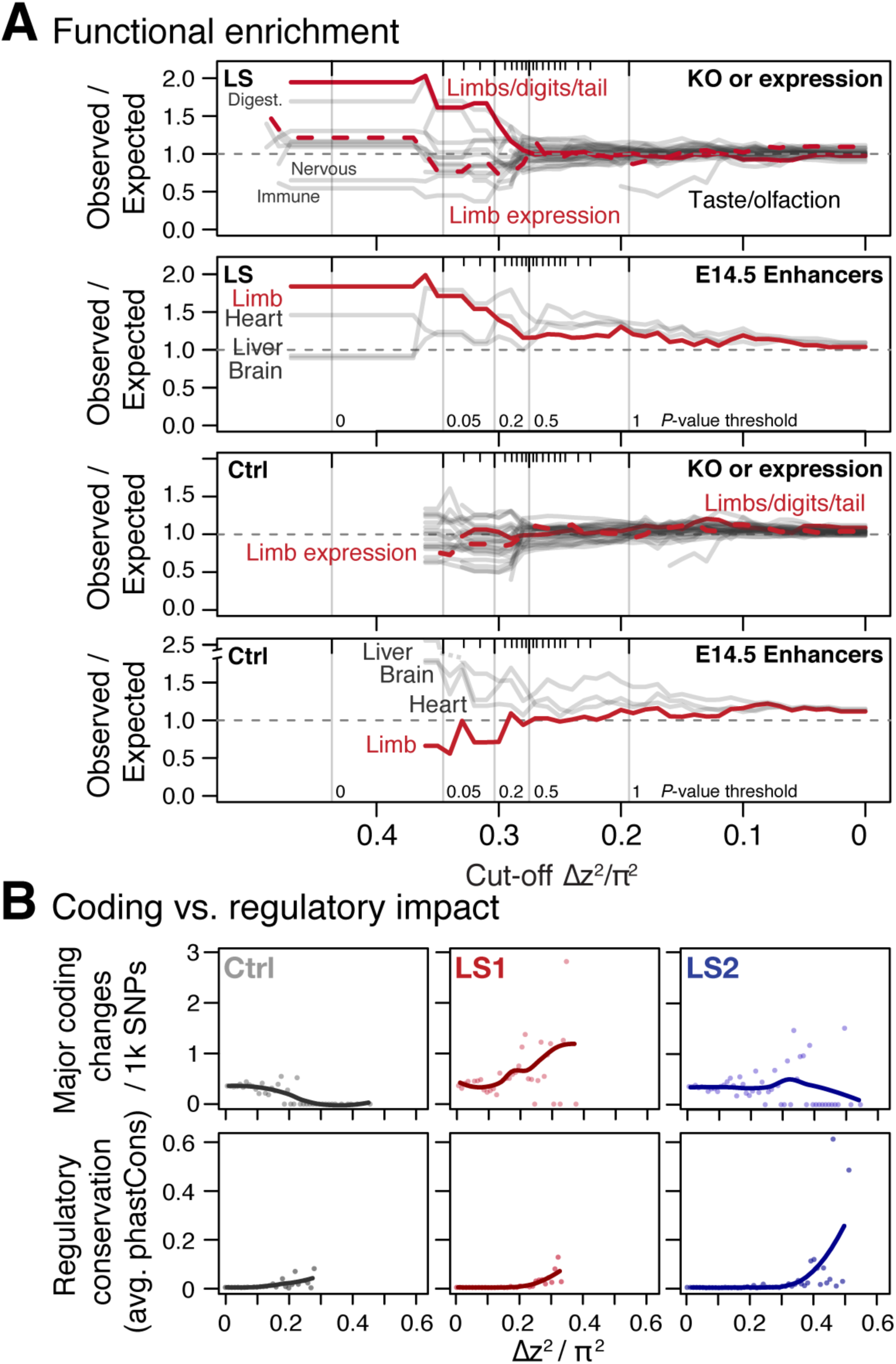
Loci associated with selection response in Longshanks lines show enrichment for limb function likely associated with ***cis*-acting mechanisms.** (**A**) Gene set enrichment analysis of knock-out phenotypes (KO) showed that selection response (here shown as Δz^2^ cut-off values, see Supplementary Methods for details on cut-off values and inclusion criteria) were found among TADs containing limb and tail developmental genes (red solid lines) or genes with limb expression (red dotted lines) in LS lines (top) but not in Ctrl (bottom). Among KO phenotypes, limb defects show the greatest excess out of 28 phenotypic categories (other grey lines, with other extreme categories labeled, the “normal” category is excluded here). Among developmental enhancers for limb, heart, liver and brain tissue, we also observed an association with Δz^2^ peaks in LS lines (top) for limb but not in Ctrl lines (bottom). The simulated significance thresholds based on *H*_*INF,* *max* *LD*_ are also shown for reference (vertical grey lines). The data from the LS line suggest that enrichment start to increase around *P* ≤ 0.5 threshold and remained largely stable by *P* ≤ 0.05, corresponding a cut-off of around 0.33 π^2^. (**B**) Coding vs. regulatory impact. Frequency of moderate to major coding changes (top panels, amino acid changes, frame-shifts or stop codons), or average conservation score of regulatory sequences immediately flanking SNPs (based on conservation among 60 eutherian mammals; bottom panels) were used as proxies to estimate the functional impact of coding and regulatory mutations, respectively. In LS1, major coding changes became more common at high Δz^2^ ranges; however the rate of SNPs with potentially major phenotype consequences did not increase in LS2 and in fact seems to decrease in Ctrl. In contrast, regulatory changes showed increased conservation associated with greater allele frequency shifts or Δz^2^ in all three lines, except that SNPs with large shifts and strong conservation were more abundant in LS1 and LS2. Trend lines are shown with LOESS regression but statistical comparisons were performed using linear regressions.

**fig. S8.**
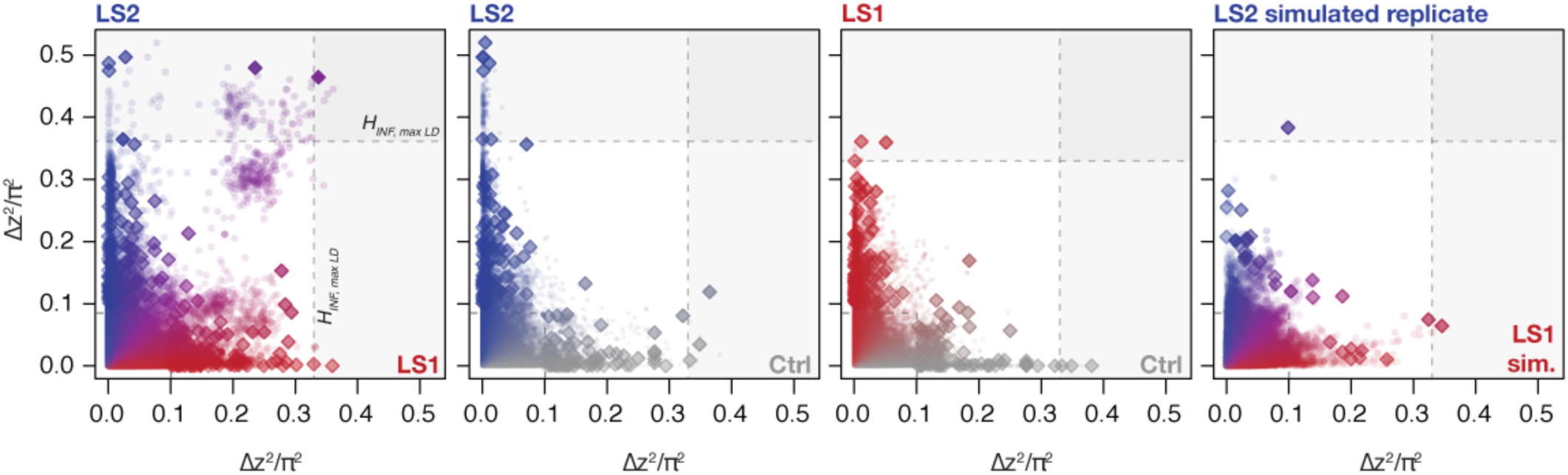
Changes in Δz^2^ **across lines.** Shown are changes in Δz^2^ in individual 10 kbp windows (all windows: circles; peak windows: diamonds). Generally there were no clear differences in Δz^2^ along the axes except a slight increase in LS2. When taken as a joint LS1–LS2 comparison, however, we observed that many windows show shifts in both LS1 and LS2 (left panel). In contrast, only very few windows show parallelism in Ctrl–LS2 and Ctrl–LS1 comparisons (middle two panels). The right panel shows a single selected simulated replicate (selection pressure 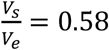 ; maximum LD) found to have among the greatest extent of parallel Δz^2^ among the replicates. The excess in parallel loci in observed results is clear both among the significant loci at *P* ≤ 0.05 under *H*_*INF,* *max* *LD*_ and highly significant at the more relaxed *H*_*INF,* *no* *LD*_ threshold.

**fig. S9.**
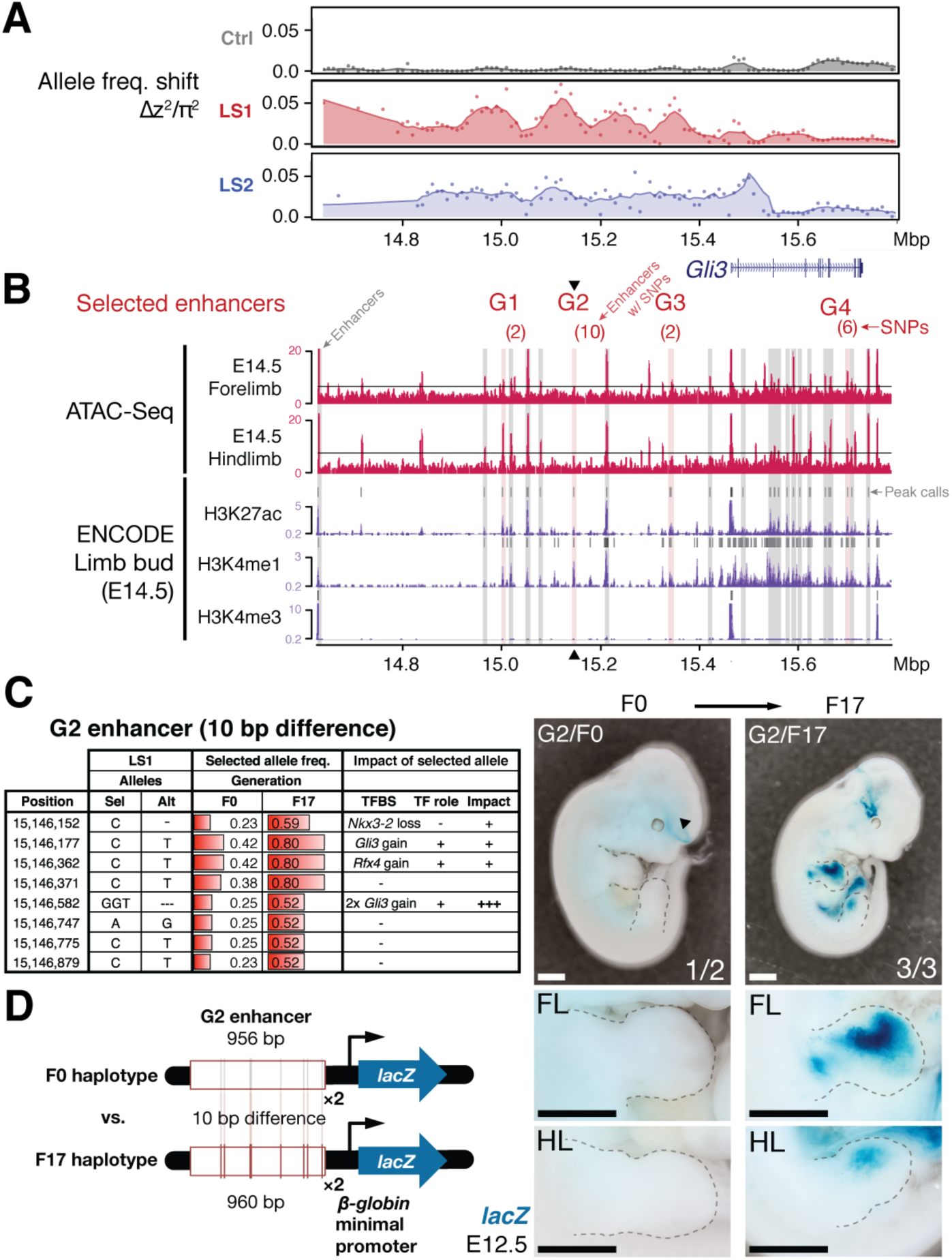
An enhancer under selection in LS1 boosts ***Gli3* expression during limb bud development.** (**A**) LS1 showed elevated Δz^2^ (peak windows: arrows) in the intergenic region in a topologically associated domain (TAD, grey shaded boxes) containing *Gli3*. (**B**) Putative limb enhancers (grey bars) were identified through peaks from ATAC-Seq (top) and histone modifications (bottom tracks, data from ENCODE project). Four of the enhancers contain mutations (in brackets) with significant allele frequency shifts between F0 and F17 in LS1 (red shading). One of the enhancers located close to the peak Δz^2^ signal (G2, arrowhead) containing 10 bp differences was chosen for transgenic reporter assay. (**C**) Analysis of individual mutations showed an average increase of 0.33 in allele frequency, with 6 mutated positions affecting predicted binding of transcription factors in the *Gli3* pathway (including 3 additional copies of *Gli3* binding site), all of which are predicted to boost the G2 enhancer activity. (**D**) The F17 G2 enhancer variants drove robust and consistent *lacZ* reporter gene expression at E12.5, recapitulating *Gli3* expression in the developing fore-and hindlimb buds (right; fig. S9). Substitution of 10 positions (F0 haplotype) led to little observable expression in the limb buds (left). Scale bars: 1 mm for both magnifications. These G2 enhancer gain-of-function mutations (contrasting the major allele between F0 and F17) may confer an advantage under selection for increased tibia length. Scale bar: 1 mm for both magnifications.

**fig. S10.**
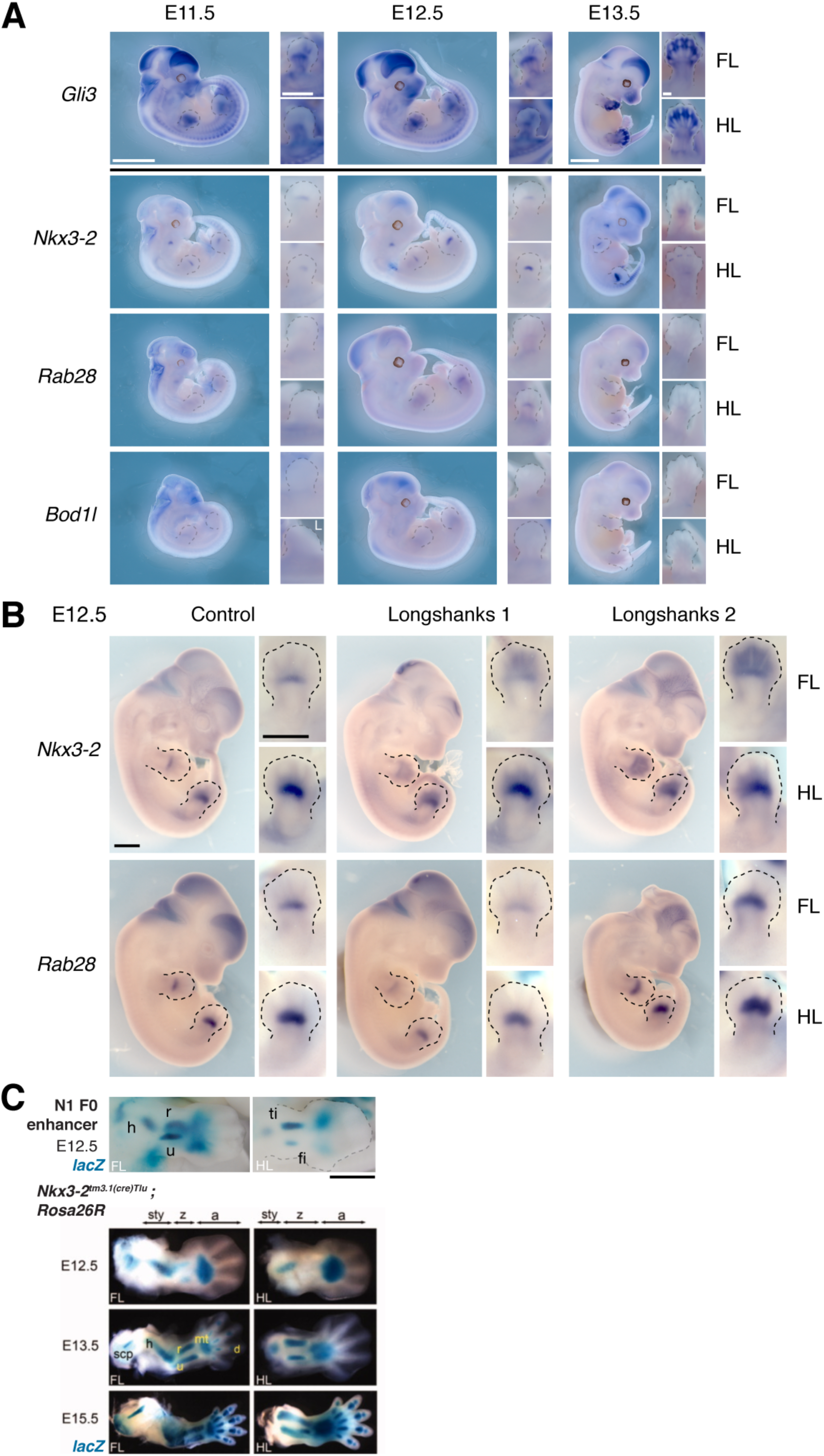
Gene expression patterns at the ***Gli3* and *Nkx3-2* candidate intervals.** (**A**) *Gli3* expression was determined using *in situ* hybridization. *Gli3* was robustly expressed during limb development in both developing fore-and hindlimb buds, especially in the autopod (hand/foot plate). Lower panel shows expression of *Nkx3-2* and its neighboring genes *Rab28* and *Bod1l*. The stronger expression of *Nkx3-2* in the developing limb buds as well as the known role of *Nkx3-2* in bone maturation (*30*) strongly argues for *Nkx3-2* being the gene underlying the selection response at the Chr5 locus. Scale bars: 1 mm for whole-mounts; 0.5 mm for limb buds. “L” indicates left side. (**B**) We collected E12.5 embryos from each line and performed *in situ* hybridization to determine the sites and level of expression of *Nkx3-2* and *Rab28* in the Longshanks (right columns) and Control (left column) lines. Both genes are expressed in similar sites overall and specifically in the developing fore-and hindlimb buds in the region of the presumptive zeugopods. These data indicate common sites of expression and rule out qualitative presence/absence differences in expression. (**C**) Although the N1 enhancer pattern appear to differ from endogenous *Nkx3-2* expression, it matches the pattern of *Nkx3-2* expression, as indicated in (*30*). The use of a *Nkx3-2 Cre-*driver line suggested possibly undetected early expression of *Nkx3-2* prior to bone formation in the limb buds (lineage tracing experiment using a *Cre*-driver and revealed through crossing to *Rosa26R*, a *lacZ* reporter line). Image modified from (*30*), reused with permission. Scale bar: 1mm. h, humerus; r, radius; u, ulna; ti, tibia; fi, fibula; sty, stylopod; z, zeugopod; a, autopod; scp, scapula; mt, metacarpals; d, digits.

**fig. S11.**
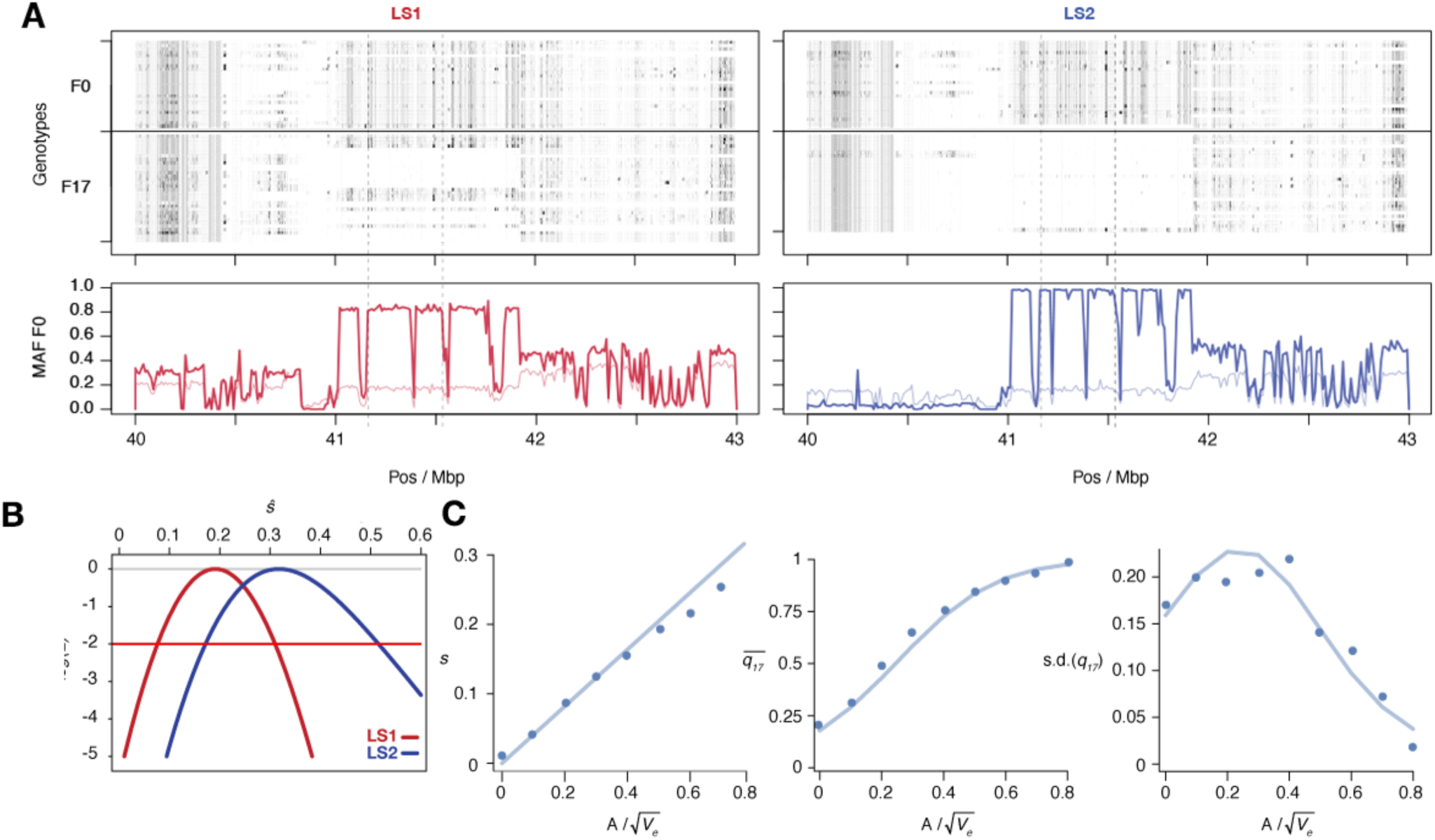
Selection at the ***Nkx3-2* locus.** (**A**) Raw genotypes from the F0 and F17 generations from LS1 (left) and LS2 (right) are shown, clearly indicating the area under the selective sweep. The genotype classes are shown as C57BL/6J homozygous (BL6, light grey), heterozygous (black) and alternate homozygous (dark grey). Lower Panel: Tracking MAF from both lines show that the originally rare F0 allele (thin line) rose to high frequency at F17 (thick lines). The plateau profile from both lines suggested that an originally rare allele became very common by F17 in both lines (see raw genotypes). Note that in LS2 F17 the region is fixed for the BL6 allele except the bottom-most individual). (**B**) The log likelihood of the selection coefficient, *s*, for LS1, LS2 (red, blue), based on transition probabilities for a Wright–Fisher population with the appropriate *N*_*e*_. The horizontal red line shows a loss of log likelihood of 2 units, which sets conventional 2-unit support limits. (**C**) Simulations of an additive allele with effect *A* on the trait; 40 replicates for each value of *A*. Left: The selection coefficient, estimated from the change in mean allele frequency, as a function of 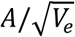; the line shows the least-squares fit 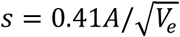. Middle: dots show the mean allele frequency at generation 17; the line shows the prediction from the single-locus Wright– Fisher model, given 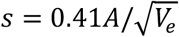. Right: the same, but for the standard deviation of allele frequency.

